# Transcriptional circuitry of NKX2-1 and SOX1 defines an unrecognized lineage subtype of small cell lung cancer

**DOI:** 10.1101/2022.03.06.483161

**Authors:** Ranran Kong, Ayushi S. Patel, Takashi Sato, Seungyeul Yoo, Li Bao, Abhilasha Sinha, Feng Jiang, Yang Tian, Maya Fridrikh, Shuhui Liu, Jie Feng, Xijing He, Jiantao Jiang, Yuefeng Ma, Karina Grullon, Dawei Yang, Charles A. Powell, Mary Beth Beasley, Jun Zhu, Eric L. Snyder, Shaomin Li, Hideo Watanabe

## Abstract

**Rationale:** The current molecular classification of small cell lung cancer (SCLC) based on expression of four lineage transcription factors still leaves its major subtype SCLC-A as a heterogeneous group, necessitating more precise characterization of lineage subclasses.

**Objectives:** To refine the current SCLC classification with epigenomic profiles and to identify features of the re-defined SCLC subtypes.

**Methods:** We performed unsupervised clustering of epigenomic profiles on 25 SCLC cell lines. Functional significance was evaluated by cell growth, apoptosis and xenograft using CRISPR-Cas9-mediated deletion. The specific cistromic profiles by ChIP-seq and its functional transcriptional partners using co-immunoprecipitation followed by mass spectrometry were determined. *Rb1^fl/fl^Trp53^fl/fl^* and *Rb1^fl/fl^Nkx2-1^fl/fl^* mouse models were engineered to explore the function of *Nkx2-1* in tumor initiation and differentiation. H3K27ac profiles were analyzed to reveal 6 human SCLC specimen and 20 mice tumors epigenomic landscapes.

**Measurements and Main Results:** We identified an epigenomic subclusters of the major SCLC-A subtype, named SCLC-Aα and SCLC-Aσ. SCLC-Aα was characterized by the presence of a super-enhancer at the *NKX2-1* locus, which was observed in human SCLC specimens and a murine SCLC model. We found NKX2-1, a dual lung and neural lineage factor, is uniquely relevant in SCLC-Aα. We further found maintenance of this neural identity in SCLC-Aα is mediated by collaborative transcriptional activity with another neuronal transcriptional factor SOX1.

**Conclusions:** We comprehensively describe an additional epigenomic heterogeneity of the major SCLC-A subtype, and define SCLC-Aα subtype by the core regulatory circuitry representing NKX2-1 and SOX1 super-enhancers and their functional collaborations to maintain neuronal linage state.

## INTRODUCTION

Small cell lung cancer (SCLC) is an aggressive, lethal, and rapidly progressing neuroendocrine (NE) malignancy, accounting for ~15% of lung cancer incidence and 250,000 deaths worldwide annually (1). The five-year survival rate for patients with SCLC is only 6% compared to 19% for non-SCLC. The lack of progress in SCLC treatment is largely attributable to the inadequate molecular understanding. Advances in genomic technology have enabled new molecular characterization of SCLCs. Genome-wide methylation and transcriptomes of human SCLCs and PDX models identified NE and non-NE subtypes. NE subtype is defined by the expression of neuronal lineage transcription factors (TFs) ASCL1 and/or NeuroD1 (2). Two recent studies described the non-NE SCLCs, one signified by YAP1 expression and another by POU2F3 expression (3, 4). In light of these findings, a consortium proposed a new molecular classification of SCLCs that distinguishes them into four subtypes based on the expression of four lineage-defining transcription factors: ASCL1, NeuroD1, YAP1 and POU2F3 referred to as SCLC-A, SCLC-N, SCLC-Y and SCLC-P, respectively (5, 6). The classification, while marking significant advance, is still a work in progress. After that report, a newer study proposed an inflamed SCLC subtype (SCLC-I) (7). In addition, not all SCLCs clearly fall into the four clusters as they often express multiple factors at variable levels, especially in the SCLC-A subtype which account for 70% of SCLCs (8). This wide range of heterogeneity warrants novel approaches to develop a more precise classification of SCLC based on lineage subtype.

Lineage-specific transcription factors and the presence of SE at their gene locus are critical determinants of cellular identity (9). In our study, we applied epigenomic approaches to investigate the differential enhancer landscapes in SCLC cell lines, human SCLC samples and autochthonous murine SCLCs from a genetic engineered mouse model (GEMM) to reveal additional intertumoral heterogeneity, providing comprehensive evidence of distinct histone modification profiles. Furthermore, we determined a novel mechanism of subtype-specific transcriptional regulation in SCLC, integrating the core transcriptional networks and their collaborative regulation of NE target genes.

## METHODS

For a detailed description of the methods, please see the online supplement.

### Cell lines

Cell lines used in this study are listed in Table E1.

### Chromatin Immunoprecipitation followed by sequencing (ChIP-seq)

ChIP-seq was performed as described previously (10). Protein G magnetic beads (Dynabeads: Life Technologies) were pre-incubated with anti-H3K27ac (Abcam, ab4729) or anti-NKX2-1 (Bethyl Laboratories, A300-BL4000) or anti-SOX1 (R&D, AF3369) antibodies. Up to 10 ng of DNA was used for the library construction using NEBNext Ultra II DNA Library Prep Kit (NEB, E7645). Sequencing was performed on NextSeq 500 (Illumina).

### RNA-seq

The procedure was performed as previously described (8, 10).

### Lentiviral transduction of genes and CRISPR-Cas9 genome editing

*FLAG-NKX2-1* or *FLAG-GFP* open reading frame (ORF) was cloned into pLEX_306 (a gift from David Root, Addgene plasmid #41391). Cells stably expressing Cas9 were generated by infection with the lentiCas9-Blast plasmid (Addgene # 52962, a gift from Feng Zhang). sgRNAs targeting *NKX2-1* or *SOX1* were selected from Brunello library (Table E2) (11). Additional details are provided in the online supplement.

### Immunohistochemistry (IHC)

Immunohistochemical analyses were performed on xenograft tumor specimens, human primary SCLC tumor specimens and genetic engineered mouse model (GEMM) tumor specimens with anti-NKX2-1 (1:100; Santa Cruz, sc-53136), anti-SOX1 (R&D, AF3369) or anti-ASCL1 (1:100, BD, 556604) antibody and detected with the Vectastain ABC kit.

### Co-immunoprecipitation and LC-MS/MS

Chromatin fraction was prepared and co-immunoprecipitated with the methods described previously (12) with modifications. Peptides were detected, isolated, and fragmented to produce a tandem mass-spectrum of specific fragment ions for each peptide with LTQ Orbitrap Velos Pro ion-trap mass spectrometer (Thermo Fisher Scientific, Waltham, MA).

### Cell proliferation assay

AlamarBlue Cell Viability Reagent was used to analyze cell proliferation. Cells were plated at a density of 3,000 cells/well with five replicates in a 96-well plate.

### Xenograft model

The procedure was performed as previously described (ref). All xenograft studies were approved by the institutional Animal Care and Use Committee at Icahn School of Medicine at Mount Sinai (IACUC-2018-0021).

### Annexin-V staining

Alexa Fluor^®^ 488 annexin V/Dead Cell Apoptosis Kit (Thermo Fisher Scientific) was used according to manufacturers’ recommendations. Fluorescence was measured by FACS Canto II (BD Bioscience).

### In situ proximity ligation assay (in situ PLA)

*In situ* PLA was performed with DuoLink in Situ Reagents (Sigma) according to the instructions. Images were acquired with a ZEISS Axiocam 503 mono confocal microscope.

### Mice and tumor initiation

Mice harboring Rb1^flox^, Trp53^flox^ and Nkx2-1^flox^ alleles have been previously described (13, 14). Tumors were generated by intratracheal delivery of Ad5CMV-Cre adenovirus (University of Iowa, Gene Transfer Vector Core) as described (15). All studies were approved by the Committee for Animal Care at MIT (A-3125-01).

### Data analysis

Details are provided in the online supplement.

### Statistical Analysis

Details of specific analyses are provided in the figure legends.

### Data and materials availability

SCLC cell lines, human primary SCLC tumors, RP and RPN tumors H3K27ac ChIP-seq, Cell lines and RP and RPN tumors RNA-seq, NKX2-1 and SOX1 ChIP-seq are deposited in NCBI GEO: GSE183373.

## RESULTS

### Epigenomic profiling subdivided SCLC-A by differential super-enhancers

Genome-scale chromatin organization of the cell more faithfully reflects their lineage state than transcriptomics (16). To refine current SCLC classification, we profiled genome-wide H3K27ac to define SE regions across 16 SCLC cell lines. Unsupervised hierarchical clustering on SEs on genes for transcriptional regulation with additional public data (4) revealed four subclasses (Figure 1A), supported by principal component analysis (PCA) (see Figure E1A in the online data supplement). Our clusters recapitulated aspects of the expression classification. Cluster III corresponds to SCLC-N (Figure 1A; purple). Cluster IV represents SCLC-P and SCLC-Y (Figure 1A; blue). Notably, epigenomic clustering distinguished SCLC-A into two distinct Clusters I and II (Figure 1A; red and green) referred to as SCLC-Aα and SCLC-Aσ, while *ASCL1* SE is present in 92.9% (13/14) of SCLC-A (Figure E1B).

**Figure 1.**
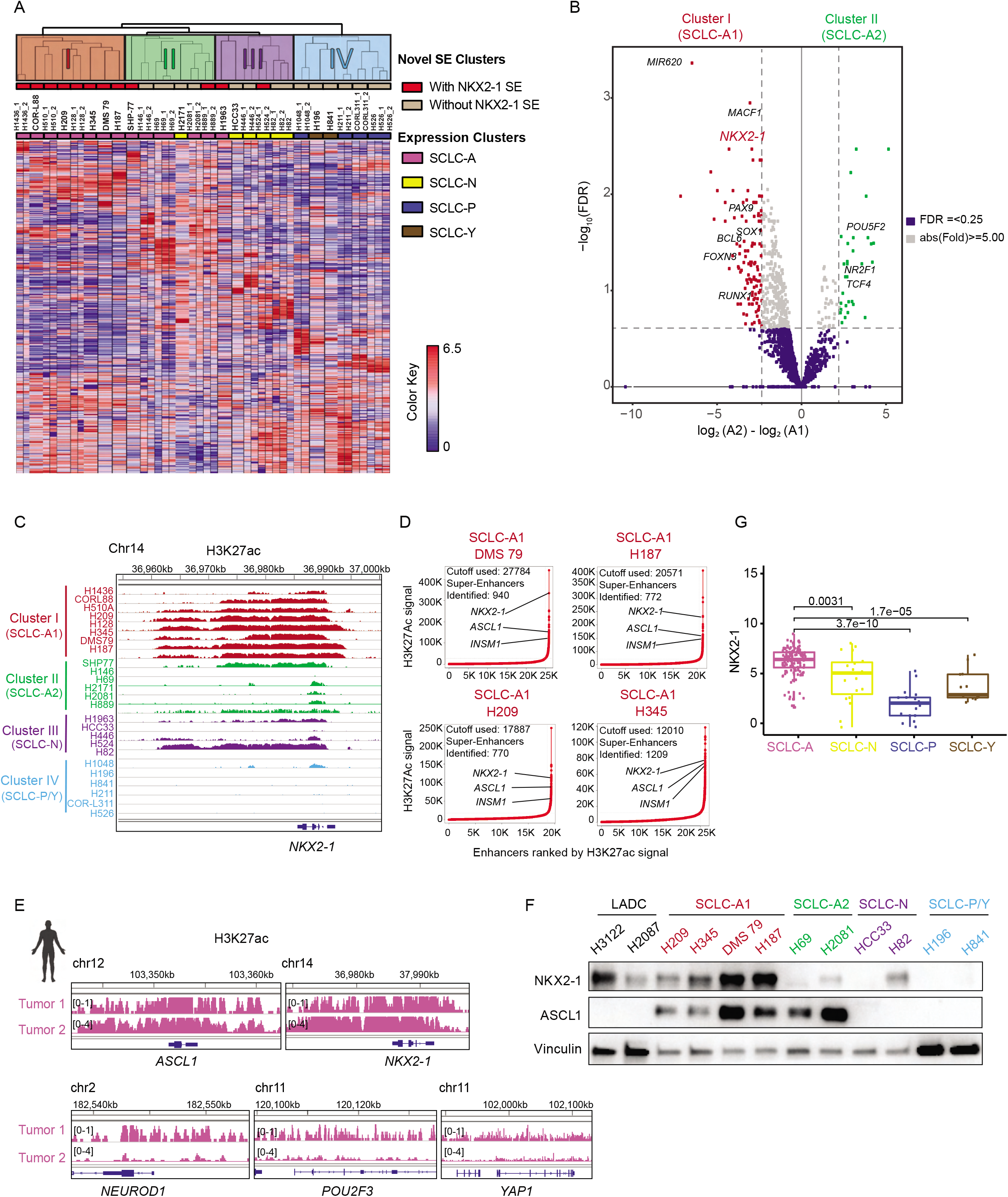
Epigenomic profiling subdivided SCLC-A by differential super-enhancers. (**A**). Unsupervised hierarchical clustering of 25 SCLC cell lines using SE signals near transcriptional regulator genes. The color scale indicates the intensity of SE signals normalized to linear scores using the sum of squares of the values. Each row represents a SE locus (747 loci). The color shades on the dendrogram indicate four novel SE clusters, the top red columns display the presence of SE near the *NKX2-1* locus, and the bottom column colors denote the consensus expression clusters. (**B**). Volcano plot showing 140 differentially enriched SEs (110 enriched in SCLC-Aα and 30 enriched in SCLC-Aσ) at thresholds of absolute fold change ≥ 5 and FDR ≤ 0.25 depicted as dotted lines. (**C**). Genome view tracks of H3K27ac signal at the *NKX2-1* locus in four clusters of SCLC cell lines. Normalized scales are shown in Figure E1B. (**D**). SE plots with all enhancers rank-ordered by H3K27ac signals in DMS 79, NCI-H187, NCI-H209 and NCI-H345. Dotted lines depict tangential cutoffs to define SEs. (**E**). Genome view tracks of H3K27ac signals at the *ASCL1, NEUROD1, POU2F3, YAP1* and *NKX2-1* loci in 2 human SCLC specimens of SCLC-A subtype. (**F**). Protein expression of NKX2-1, ASCL1 and vinculin as a loading control in SCLC cell lines from different clusters. *NKX2-1*-amplified adenocarcinoma cell lines NCI-H3122 and NCI-H2087 are shown as reference. (**G**). Box and jitter plots showing the distribution of *NKX2-1* mRNA expression in SCLC-A, SCLC-N, SCLC-P, and SCLC-Y subtypes of 135 primary SCLC tumors.

To understand the difference underpinning SCLC-Aα and SCLC-Aσ, we identified 140 differentially enriched SEs. Among 110 SEs enriched in SCLC-Aα was *NKX2-1* (Figure 1B), where H3K27ac signal was largely absent in other subclasses (Figure 1C). The 30 SEs enriched in SCLC-Aσ included *TCF4,* a transcription factor that regulates synaptic plasticity (17) (Figure 1B and Figure E1B). NKX2-1, also known as *TTF1,* a lineage transcription factor essential for the development and maintenance of lungs, brain, and thyroid, is a frequently amplified lineagespecific oncogene and a well-known marker to diagnose lung adenocarcinoma (LADC) (18) (19, 20). However, *NKX2-1* is not significantly amplified in human SCLCs (21) nor in the SCLC lines used (Figure E1C). Nonetheless, SCLC-Aα cell lines harbor *NKX2-1* SE, whereas SCLC-Aσ cell lines do not (Figure 1D and Figure E1D). Local H3K27ac enrichment pattern at the *NKX2-1* locus in SCLC-Aα was almost identical to LADCs with *NKX2-1* amplification (Figure E1E). In 6 human SCLC specimens, we found one SCLC-P tumor with *POU2F3* SE, and threes mixed SCLC-A/N tumors with SEs near *ASCL1* and *NEUROD1* loci (Figure E1F). Notably, we observed two tumors representing SCLC-Aα with high signals at *ASCL1* and *NKX2-1* loci (Figure 1E).

By immunohistochemistry (IHC) on another 34 human SCLC cohort, 65% (22/34) of cases were NKX2-1-positive (Figure E2A), consistent with prior studies (22–26). In cell lines, SCLC-Aα expressed substantially higher levels of NKX2-1 protein than other SCLC subtypes and equivalent to *NKX2-1*-amplified LADCs (Figure 1F). Furthermore, RNA-seq datasets (5, 27) confirmed *NKX2-1* is highly expressed in SCLC (Figure E2B), particularly in SCLC-A (Figure 1G), comparable to the levels in LADC (Figure E2C), consistent with a previous report (22). Nonetheless, non-bimodal distribution of *NKX2-1* expression suggests that the distinction between SCLC-Aα and Aσ is not discernible from expression analysis, perhaps analogous to the findings that *NKX2-1* is almost universally expressed in LADC yet only amplified in 10-15%, which we previously characterized as a distinct biological subgroup (19, 28).

### *NKX2-1* survival dependency is unique to SCLC-Aα

A recent study implicated that NKX2-1 promotes SCLC growth and regulates NE differentiation and anti-apoptosis (29). We deleted *NKX2-1* in SCLC-Aα cell lines (Figure 2A) and found it significantly suppressed cell growth and increased annexin-V-positive populations (Figure 2C and E3A). By contrast, ablating NKX2-1 in SCLC cells from other clusters (Figure 2B) had little effect. Given that acute removal of a master lineage factor in established cancer often leads to a halt in cell growth (30), these data suggest NKX2-1 does not play a general oncogenic role in all lineage states but its function is uniquely relevant to SCLC-Aα.

**Figure 2.**
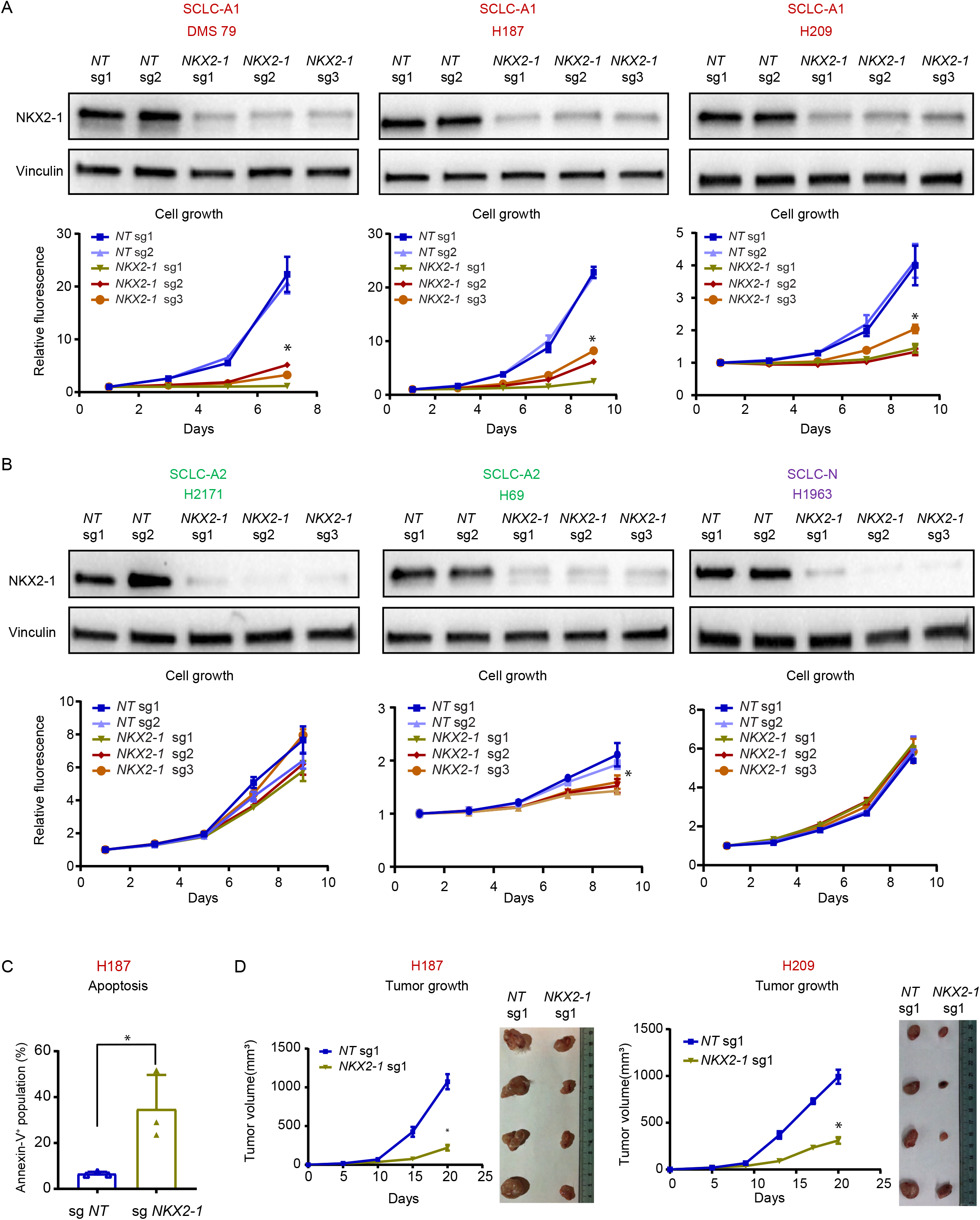
*NKX2-1* survival dependency is unique to SCLC-Aα. (**A**). *(Top)* Immunoblots showing protein expression of NKX2-1 and vinculin as a loading control in SCLC-Aα cell lines with three *NKX2-1* sgRNAs, or two non-target (NT) sgRNAs. *(Bottom)* Cell growth curve for SCLC-Aα cell lines infected with *NKX2-1* sgRNAs or NT sgRNAs. Data are shown as mean ± SD (n=3); *P < 0.0001 by two-sided two-way ANOVA. (**B**). *(Top)* Immunoblots showing protein expression of NKX2-1 and vinculin as a loading control in SCLC-Aσ (NCI-H2171, NCI-H69) and SCLC-N (NCI-H1963) cell lines with *NKX2-1* sgRNAs, or nontarget (NT) sgRNAs. *(Bottom)* Cell growth curve for cell lines infected with *NKX2-1* sgRNA or NT sgRNAs. Data are shown as mean ± SD (n=3); *P < 0.0001 by two-sided two-way ANOVA. (**C**). Bar plots showing percentage of apoptotic NCI-H187 cells infected with NT sgRNA and *NKX2-1* sgRNA, measured by annexin V staining. Means ± SD of three biological replicates. *P < 0.0001, t test. Representative data for NT sg1 and NKX2-1 sg1 can be found in Figure E2A. (**D**). *(Left)* Tumor growth curve for xenograft tumors of NCI-H187 cells infected with *NKX2-1* sg1 or NT sg1 *in vivo.* Mean ± SD of four biological replicates. *P < 0.0001, t test. Xenograft tumors were resected at 3 weeks after inoculation. *(Right)* Images of resected xenograft tumors of NCI-H209 cells infected with NT sg1 or *NKX2-1* sg1 *in vivo* at day 23.

We further found a significant decrease in tumor volume for NKX2-1-deleted SCLC-Aα cells in xenograft (Figure 2D and Figure E3B), confirming its essentiality *in vivo.* No obvious histological differences were observed in the cells that formed small tumors which escaped from NKX2-1 deletion (Figure E3C). In a short period after NKX2-1 deletion *in vitro,* expression of NE markers was not significantly altered (Figure E3D). Overall, these data suggest that NKX2-1 is functionally critical for SCLC-Aα *in vitro* and *in vivo.*

### Genomic occupancy of *NKX2-1* in SCLC-Aα cells distinct from that in LADC cells

In LADC, NKX2-1 regulates gene expression and shares its cistromes with normal lung (19); however, SCLCs typically do not express canonical NKX2-1 targets. Given NKX2-1’s roles in subsets of cortical, striatal, and pallidal neurons (31), we speculated its potential role for neuronal differentiation in SCLC-Aα. Therefore, we profiled NKX2-1 cistromes by ChIP-seq in 5 SCLC-Aα cell lines and compared them to that from 3 LADC cell lines. Pairwise correlation coefficient of all NKX2-1-bound sites and PCA revealed histology-specific binding profiles (Figure 3A and E4A). Regions 4 (low-intensity signals) and 1 (high-intensity signals) are common between SCLC-Aα and LADC (Figure 3B), which included the *RUNX1* locus (Figure 3C). Region 3 was unique to LADC that included *LMO3,* an essential mediator of NKX2-1 functions in LADCs (19). Region 2 was unique to SCLC-Aα that included *DCC,* a receptor for netrin required for axon guidance (32) (Figure 3C). These data suggest NKX2-1 regulates distinct transcriptional programs across these two lineage states.

**Figure 3.**
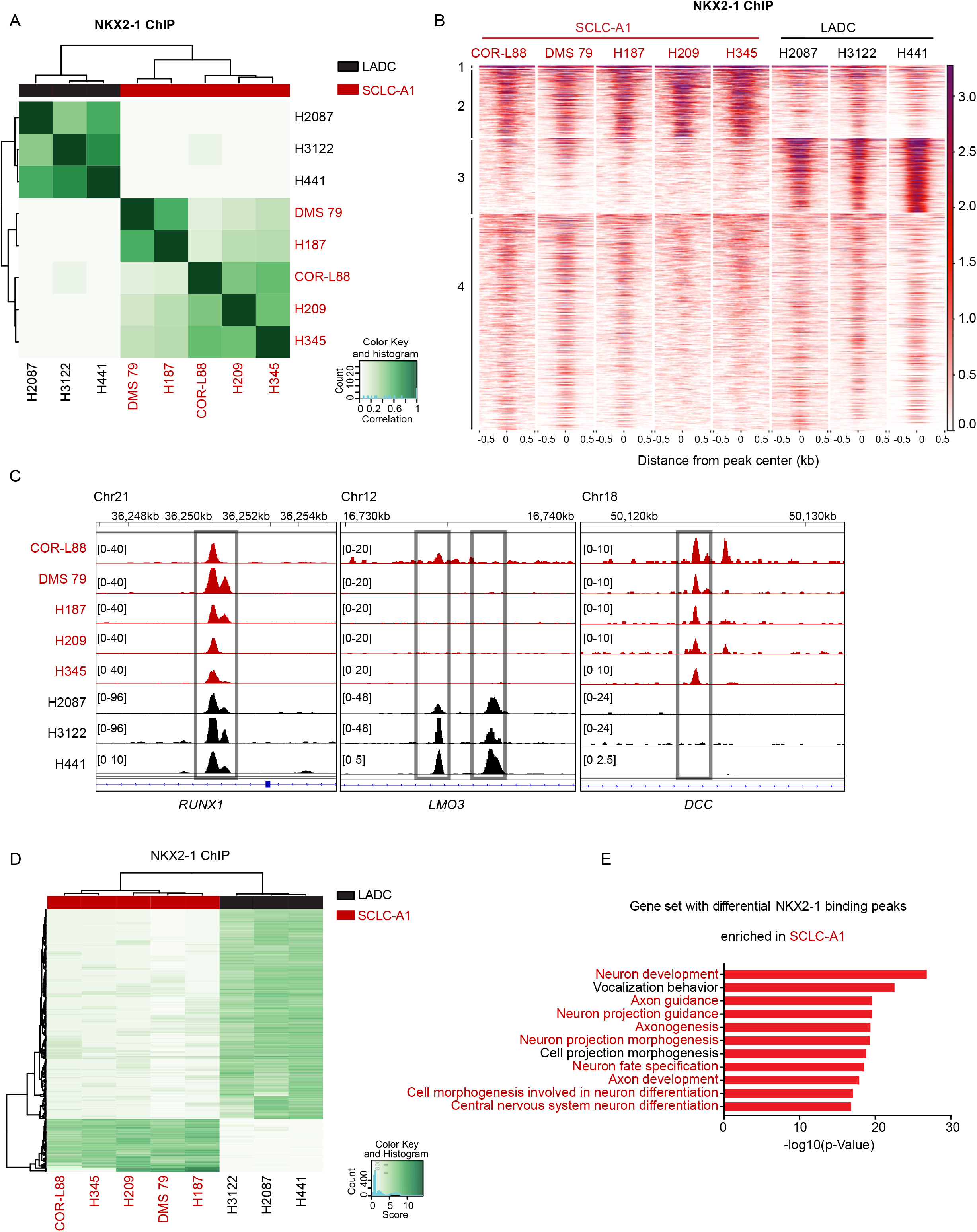
Genomic occupancy of *NKX2-1* in SCLC-Aα cells distinct from that in LADC cells. (**A**). Correlation matrix depicting pairwise comparisons of identified NKX2-1 binding peaks in the 5 SCLC-Aα cell lines and in 3 LADC cell lines. Color scale represents degree of Pearson’s correlation coefficient. (**B**). Heatmap showing signals from NKX2-1 ChIP-seq data in the 5 SCLC-Aα cell lines and 3 LUAC cell lines. ChIP signal intensity is shown by red shading. Y-axis contains all peaks that are bound by NKX2-1 in any of SCLC-Aα and LUAC (n=37,198). Peaks are divided into 4 regions by *k-means* (k=4) clustering. (**C**). NKX2-1 ChIP-seq signals in SCLC-Aα and LADC cell lines at representative loci *(RUNX1;* cluster 1, *LMO3;* cluster 3 and *DCC;* cluster 2). Red: SCLC-Aα cell lines. Black: LADC cell lines. (**D**). Heatmap showing 2,671 differentially NKX2-1 bound regions (fold change ≥ 2^5^ and FDR ≤ 1×10^-5^), 2,010 enriched in LADC and 661 enriched in SCLC-Aα, between 5 SCLC-Aα cell lines and 3 LUAC cell lines. (**E**). Enriched ontology by GREAT analyses for regions differentially bound by NKX2-1 in 5 SCLC-Aα compared to LADCs.

Ontologies on 661 SCLC-Aα-unique regions (Figure 3D and E4B) were enriched for neuronal development, differentiation, and structure (Figure 3E). By contrast, genes regulating neuronal biology were absent in LADC-unique regions (Figure E4C). Motif analysis revealed SCLC-Aα-specific sites are enriched with ASCL2, Olig2, and SOX motifs, suggesting more association with neuronal TFs (Figure E4D). In contrast, AP-1, Ets, Runx, and Tead-binding motifs are enriched in LADC-specific regions, consistent with our previous report (19). These data indicate that contribution of NKX2-1 in regulating pathways for maintenance of neural differentiation state of SCLC-Aα, distinct from its role in normal lung epithelium and LADCs.

### NKX2-1 interactome in chromatin of SCLC-Aα cells includes SOX1

NKX2-1 interacts with FOXA1 to co-regulate the expression of a set of genes for lineage control in LADC (19, 20). Therefore, to identify unique partners that interact with NKX2-1 to exert distinct transcriptional program in SCLC-Aα, we performed co-immunoprecipitation (co-IP) followed by liquid chromatography tandem mass-spectrometry (LC-MS/MS) in the chromatin fraction of two SCLC-Aα cell lines ectopically expressing FLAG-NKX2-1 (Figure 4A and Figure E5A). We found 27 overlapping NKX2-1-interacting TFs in both cell lines (Figure 4B).

**Figure 4.**
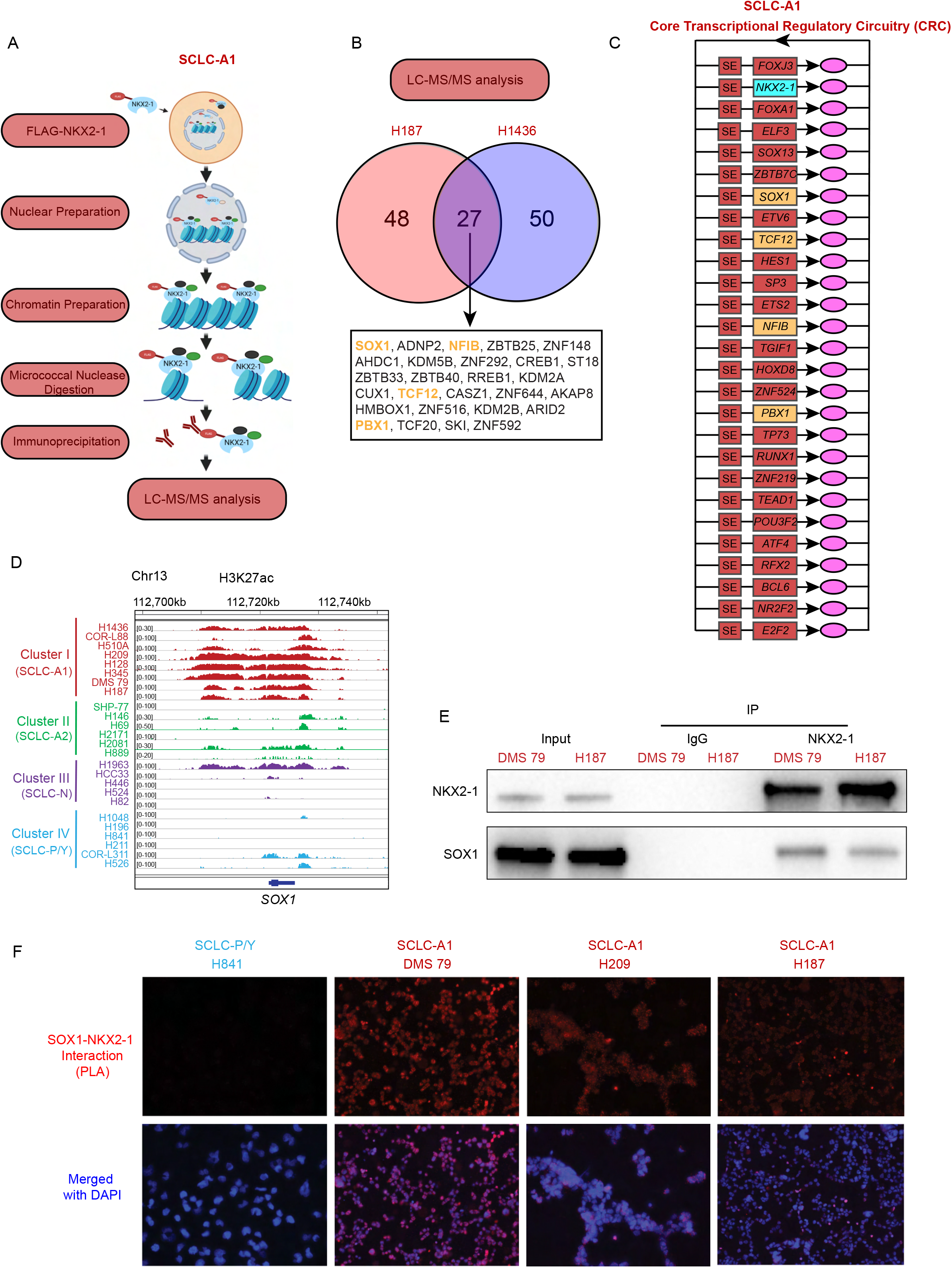
NKX2-1 interactome in chromatin of SCLC-Aα cells includes SOX1. (**A**). Experimental strategy to identify binding partners of NKX2-1 in SCLC-Aα. Chromatin prepared from the nuclear fraction of SCLC-Aα cell lines NCI-H187 and NCI-H1436 stably expressing FLAG-NKX2-1 was solubilized using micrococcal nuclease digestion. Coimmunoprecipitated proteins with anti-FLAG antibody from solubilized chromatin were identified by LC-MS/MS (see Data file S2 for the complete list). (**B**). Venn diagram showing the number of transcription factors identified in each of the 2 SCLC-Aα cell lines and the 27 overlaps found in both cell lines. The proteins indicated in yellow color were also found in CRC analysis in Figure 4C. (**C**). Core transcriptional regulatory circuitries (CRCs) commonly identified in 8 SCLC-Aα cell lines. The genes in yellow color were also found in LC-MS/MS analysis in Figure 4B. (**D**). Genome view tracks of H3K27ac signal at the *SOX1* locus in four clusters of SCLC cell lines. (**E**). Immunoblots with anti-SOX1 antibody on endogenous co-immunoprecipitates by anti-NKX2-1 antibody in SCLC-Aα (DMS 79 and NCI-H187) cells. (**F**). *In situ* proximity ligation assay signals (red fluorescence) targeting endogenous SOX1 and NKX2-1 in NCI-H841, NCI-H187, NCI-H209 and DMS 79 cells. Nuclei are counterstained with DAPI (blue). See Figure E4G for additional control.

We further identified core regulatory circuitries (CRCs) (33) for SCLC-Aα and other SCLC clusters from our H3K27ac data. Common CRCs included NEUROD1 for SCLC-N, POU2F3 and ASCL2 for SCLC-P/Y, and TCF4 in SCLC-Aσ, consistent with our differential SE analysis (Figure 1B and E5B). Common CRCs in SCLC-Aα included NKX2-1 (Figure 4C) along with SOX1, NFIB, TCF12 and PBX1 also found in the SCLC-Aα NKX2-1-interactome. Of those, H3K27ac signal at *SOX1* was highly enriched in SCLC-Aα (Figure 1B and Figure 4D), whereas *NFIB, TCF12* and *PBX1* SEs were present across SCLC subtypes (Figure E5C). *SOX1* is highly expressed in SCLC-A (Figure E5D) and positively correlated with *NKX2-1* expression in 185 SCLC tumors and cell lines (Figure E5E). SOX1 protein was also highly expressed specifically in SCLC-Aα cell lines (Figure E5F).

The interaction between NKX2-1 and SOX1 was confirmed by co-IP followed by western blot in two SCLC-Aα cell lines (Figure 4E), and further by proximity ligation assay (PLA) in 4 SCLC-Aα cell lines (Figure 4F and Figure E5G). These data suggest robust protein-protein interaction between NKX2-1 and SOX1 uniquely in SCLC-Aα.

### NKX2-1 and SOX1 co-occupy the SCLC-Aα genome to collaborate in neuronal gene regulation

Next, we profiled genome-wide binding profiles of NKX2-1 and SOX1 in four SCLC-Aα cell lines and found the majority of SOX1 peaks were bound by NKX2-1 (48.6-67.8%) (Figure 5A). The 2,652 regions co-bound by NKX2-1 and SOX1 in NCI-H187 were enriched for functions in neuron differentiation and development, consistent with the aggregate analysis (Figure E6A and E6B). We defined high-confidence NKX2-1 (5,663) and SOX1 (1,101) binding sites observed in ≧3 cell lines, and found 604 peaks (54.9% of SOX1 peaks) are overlapped (Figure 5B), which included the *NKX2-1* locus as well as neuronal genes *NRXN3* (34, 35) and *YWHAZ* (36) (Figure 5C) and ontologically enriched for functions in neuron differentiation and development (Figure 5D). These data suggest the interaction between NKX2-1 and SOX1 elicits specific cellular identity of SCLC-Aα.

**Figure 5.**
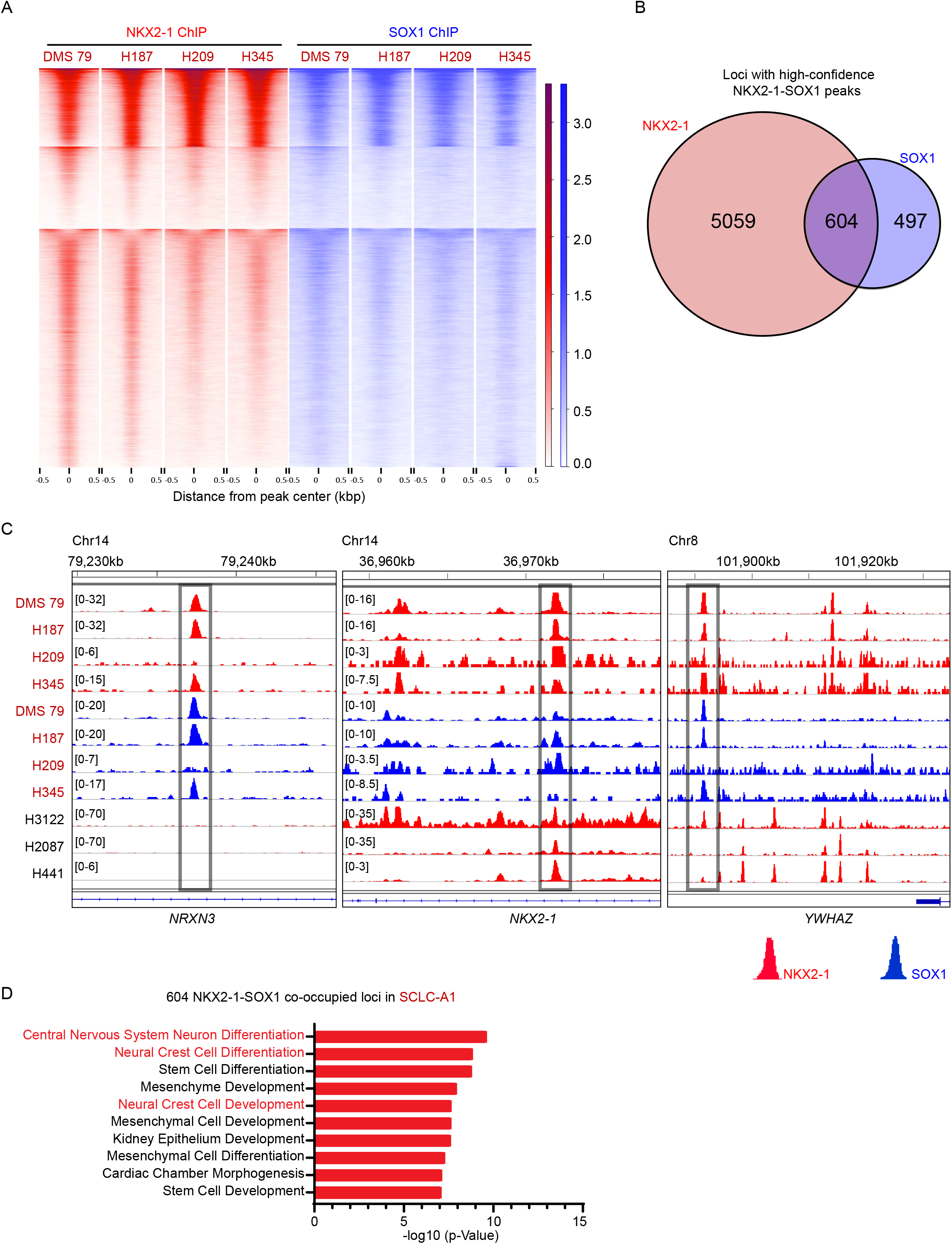
NKX2-1 and SOX1 co-occupy the SCLC-Aα genome to collaborate in gene regulation. (**A**). Heatmap showing k-means clustering (*k*=4) of signals at the common 4,051 loci from NKX2-1 (red) and SOX1 (blue) ChIP-seq data in the 4 SCLC-Aα cell lines. ChIP signal intensity is shown by shading for all NKX2-1 and SOX1 occupied loci. (**B**). Venn diagram showing the overlap of NKX2-1 common binding sites and SOX1 common binding sites present in at least 3 SCLC-Aα cell lines. (**C**). Genome view tracks of NKX2-1 (red) and SOX1 (blue) ChIP-seq signals in four SCLC-Aα (DMS 79, NCI-H187, NCI-H209 and NCI-H345) cell lines and 3 LADC cell lines at the representative NKX2-1-SOX1 co-occupied loci *(NRXN3* and *YWHAZ;* SCLC-Aα unique*, NKX2-1;* SCLC-Aα/LADC common). (**D**). Enriched ontology by GREAT analyses for regions in 604 overlapping high-confidence NKX2-1-SOX1 binding sites.

### Collaboration of NKX2-1 and SOX1 is required to control the neuronal state in SCLC-Aα

We evaluated the transcriptional effect of depletion of *SOX1* or *NKX2-1* in NCI-H187 by RNA-seq. After the abrogation of NKX2-1 or SOX1, 1,121 and 2,499 genes were significantly downregulated respectively, with 873 genes commonly downregulated, suggesting a substantial overlap in their transcriptional outputs (Figure 6A). We then queried the differentially co-downregulated genes with the 2,652 genes with transcriptional start sites (TSSs) within 50 kb of SOX1-NKX2-1 co-occupied peaks in NCI-H187, which were significantly enriched for NKX2-1 or SOX1 downregulated genes by ssGSEA analysis (Figure 6B). Leading edge analysis on these genes revealed 289 overlaps, which we found are enriched for neuron differentiation and development (Figure 6C). Together, these data suggest that NKX2-1 and SOX1 are specifically co-localized on the SCLC-Aα genome and collaboratively contribute to controlling neuronal differentiation state. We also found that SOX1 deletion, while not affecting the NKX2-1 expression, led to reduced cell growth (Figure E7A) with increased apoptosis (Figure E7B).

**Figure 6.**
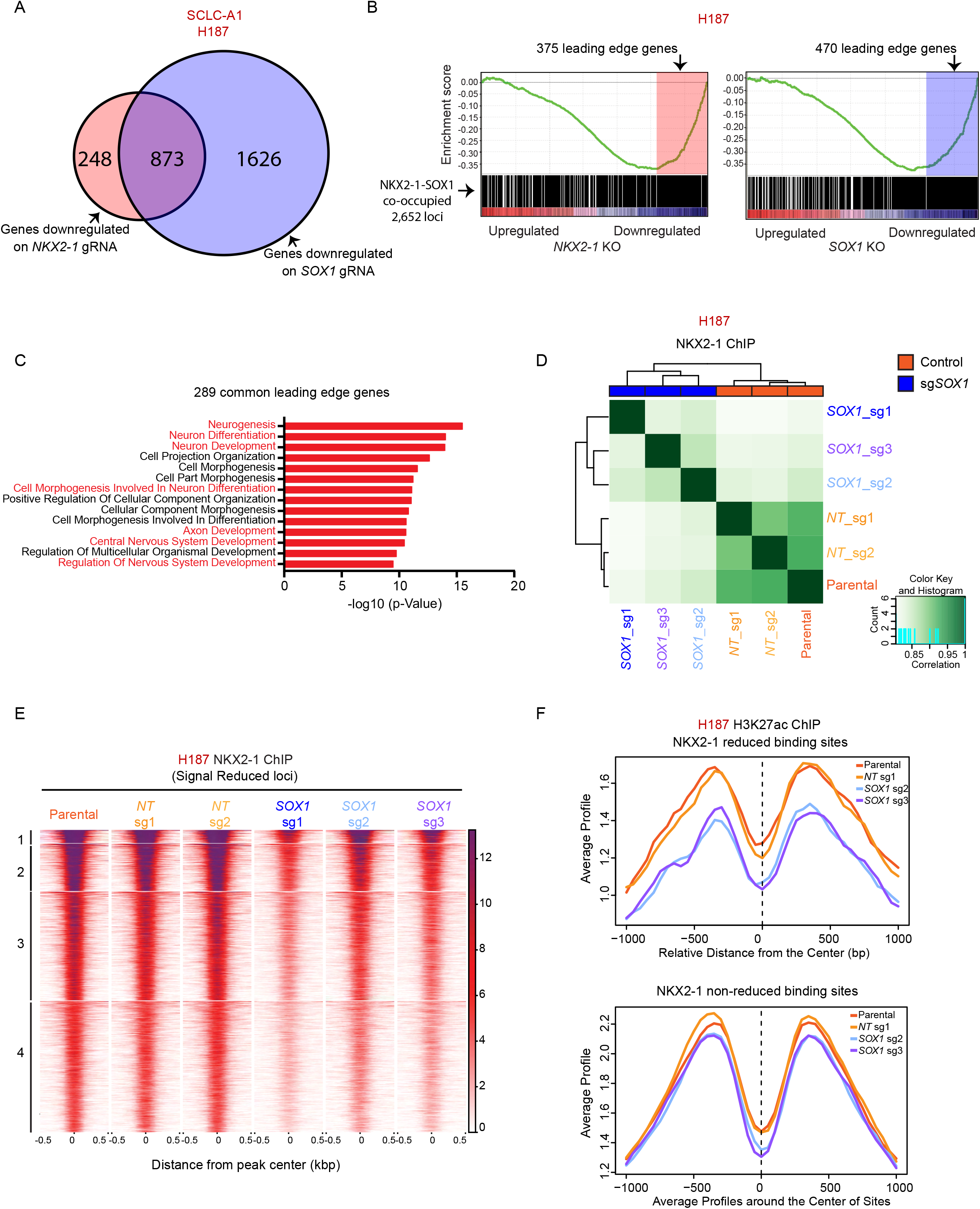
Collaboration of NKX2-1 and SOX1 is required to control the neuronal state in SCLC-Aα. (**A**). Venn diagram showing the overlap of downregulated genes after depletion of NKX2-1 (fold change > 1.4 and adjusted P < 0.05) and SOX1 (fold change > 2 and adjusted P < 0.05) in NCI-H187 cells. (**B**). Gene Set Enrichment Analysis (GSEA) of genes associated with 2,652 NKX2-1-SOX1 co-occupied loci for expression data upon repression of NKX2-1 *(left)* or SOX1 *(right)* in NCI-H187 cells, with their leading-edge genes shaded. Enrichment was significant for both (NES= −2.29, P < 0.001 for NKX2-1 and NES= −2.54, P < 0.001 for SOX1). (**C**). Enriched gene ontologies (MSigDB, biological process) for 289 common genes in leading-edge genes for both NKX2-1 and SOX1 deletions. (**D**). Correlation matrix depicting pairwise comparisons of identified NKX2-1 binding regions in three *SOX1* sgRNAs and three control cells (two non-target (NT) sgRNAs, one parental cell) of NCI-H187 cell line. Color scale represents degree of Pearson’s correlation coefficient. (**E**). Heatmap showing k-means clustering (*k*=4) of 2,067 NKX2-1 ChIP-seq loci in NCI-H187 sgSOX1 compared to control cells with reduced signal intensity. ChIP-seq signal intensity is shown by color shading. (**F**). Average H3K27ac 656 ChIP-seq signals on NKX2-1 reduced 657 binding sites and NKX2-1 non-reduced binding sites defined in Figure E6D for NCI-H187 parental, 658 NT sg1, SOX1 sg2 and SOX1 sg3 cells.

We previously found that expression of Nkx2-1 was critical for global Foxa1/2 binding pattern in the LADC genome (20). To test if SOX1 is required for NKX2-1-binding to its target genes in SCLC-Aα, we performed NKX2-1 ChIP-seq in NCI-H187 cells upon SOX1 depletion. Pairwise correlation coefficient of NKX2-1 peaks across cells deleted for SOX1 or control revealed strong correlation amongst each group (Figure 6D), supported by PCA analysis (Figure E7C). Differential analysis on NKX2-1-bound sites found 2,067 peaks decreased upon SOX1 deletion (Figure E7D). *k*-means clustering of the differential sites found relatively uniform reduction of NKX2-1 binding after SOX1 deletion (Figure 6E), where we also found decreased H3K27ac modification compared to other sites (Figure 6F), suggesting that these NKX2-1-lost sites are no longer in active state, similar to a recent finding on Sox2 in esophageal squamous cell cancer model for Klf5 (37). The data suggest that loss of SOX1 releases NKX2-1 from co-occupied loci in SCLC-Aα.

### *Nkx2-1* is essential for development of SCLC-Aα tumors in an autochthonous mouse model

The *Rb1^fl/fl^Trp53^fl/fl^* (RP) is commonly used SCLC models that recapitulates the biology of ASCL1-positive/classic human SCLCs (13) (Figure 7A). We first profiled H3K27ac to identify SEs in mSCLC tumors from RP model, where Nkx2-1 is diffusely expressed. We found all 10 tumors from 2 different mice harbored high-level H3K27ac signal around the *Ascl1* locus, but none harbored SEs at the *Neurod1, Yap1* or *Pou2f3* locus, suggesting SCLC-A subtype (Figure 7B and Figure E8A) consistent with requirement of *Ascl1* for tumor formation in this model (38, 39). We found 3 of 10 tumors simultaneously had high-level H3K27ac signals at the *Nkx2-1* and *Sox1* loci, reminiscent of SCLC-Aα, while other 7 tumors have neither. This suggests that RP can adopt an epigenetic state resembling either SCLC-Aα or SCLC-Aσ. Consistent but ambiguous expression levels by IHC suggested epigenetic profiles distinguish these lineage classes more clearly (Figure 7C).

**Figure 7.**
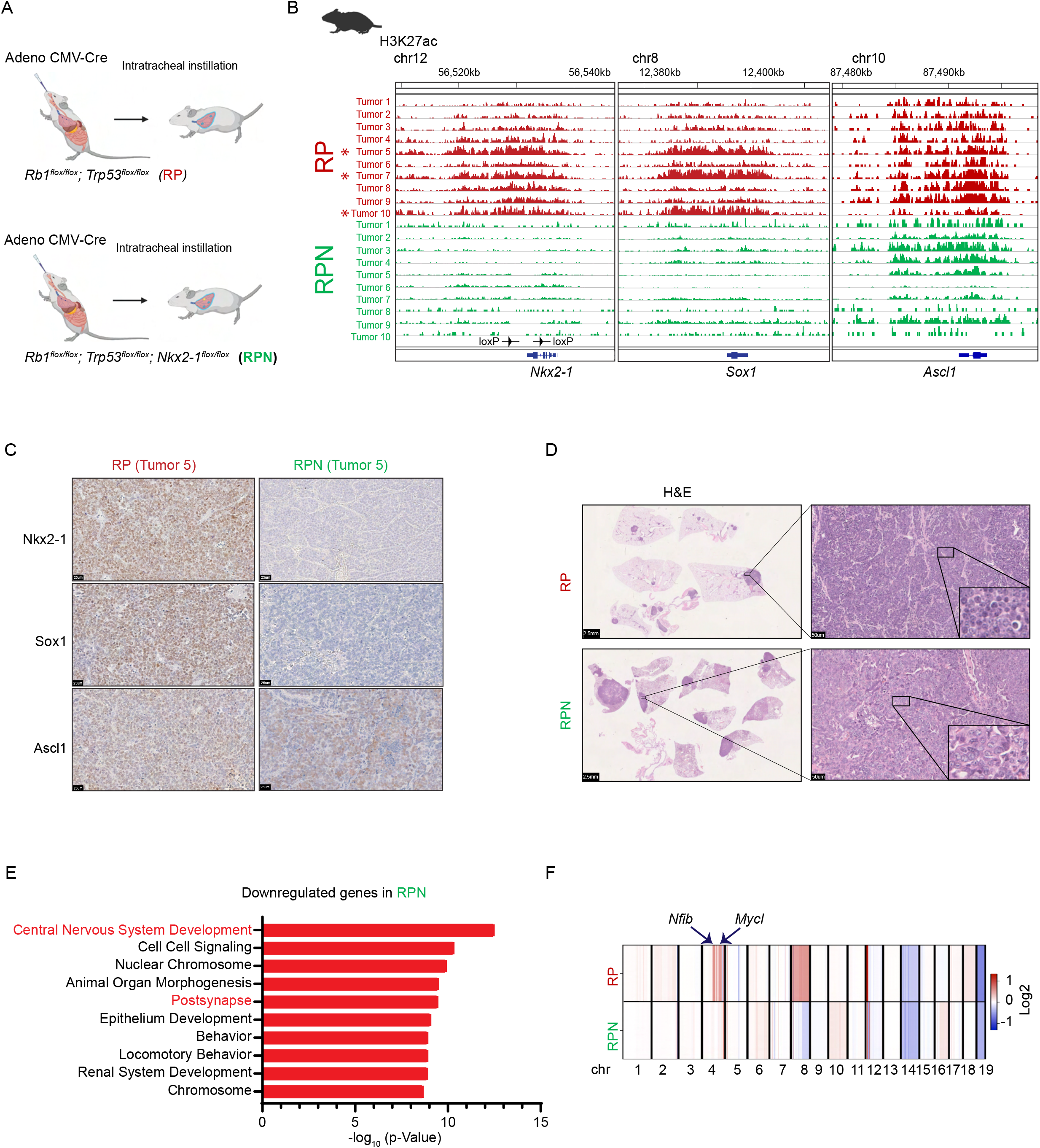
*Nkx2-1* is essential for development of SCLC-Aα tumors in an autochthonous mouse model. (**A**). Genetically engineered mouse models of SCLCs. *Rb1*^flox/flox^; *Trp53*^flox/flox^ (RP) mice or *Rb1*^flox/flox^; *Trp53*^flox/flox^, *Nkx2-1*^flox/flox^ (RPN) mice were transduced with Adeno-CMV-Cre to initiate SCLC. (**B**). Genome view tracks at the Nkx2-1, Sox1, and Ascll loci for H3K27ac ChlP-seq signals in 10 SCLC tumors from RP mice and 10 SCLC tumors from RPN mice. * indicates 3 tumors that have high-level signal at both *Nkx2-1* and *Sox1* loci. The arrowheads on *Nkx2-1* locus indicate the loxP sites in RPN mice. (**C**). Representative immune-staining images for Nkx2-1, Sox1, and Asc11 on SCLC tumors from RP and RPN mice. Scale bars, 25μm. (**D**). Representative hematoxylin and eosin (H&E) images of SCLC tumors from RP and RPN mice. Scale bars, 2.5mm and 50μm, respectively. (**E**). Enriched gene ontologies (MSigDB, biological process) for the differentially downregulated genes in RPN tumors compared to RP tumors. (**F**). Average copy number profiles in RP and RPN models. Arrows indicate the *Nfib* and *Mycl* loci that are specifically amplified in RP tumors (individual copy number profiles at those loci are available in Figure E8C).

To explore whether NKX2-1 contributes to mSCLC differentiation, we generated *Rb1^fl/fl^Trp53^fl/fl^Nkx2-1^fl/fl^* (RPN) mice. We confirmed Nkx2-1 expression was absent in their tumors, while 80% of them are Ascl1-positive suggesting that they remain representing SCLC-A. Sox1 staining was only weakly positive in 6/10 cases while 4 other cases, including the two Ascl1-negative tumors, were negative (Figure 7C). We observed no significant difference in overall survival between the two genotypes (Figure E8B). Morphologically, all but one mSCLC tumor from the RP model had classical SCLC morphology (scant cytoplasm, nuclei with fine chromatin and inconspicuous nucleoli), whereas a subset of tumors in RPN mice (15% in RPN vs. 4% in RP) exhibited a ‘non-classical’ histology defined by higher grade/vesicular nuclei by two expert pathologists (E.L.S. and M.B) (Figure 7D). These suggest that loss of *Nkx2-1* leads to stochastic propensity to form non-classical morphology, the while majority of the tumors retain classical histology representing SCLC-A.

Next, we profiled H3K27ac in 10 mSCLC tumors from four RPN mice (11-13 months post-initiation, including five classical and five non-classical). None of the tumors from the RPN mice had high-level H3K27ac signal at the *Neurod1, Yap1* or *Pou2f3* loci, whereas most of RPN tumors (7/10) harbored high-level H3K27ac signals around the *Ascl1* locus (Figure 7B and Figure E8A). Strikingly, we found signals at the *Nkx2-1* and the *Sox1* loci were absent in all RPN tumors, suggesting SCLC-Aσ subtype (Figure 7B). These data suggest that while the RP model can develop both SCLC-Aα and SCLC-Aσ, *Nkx2-1* deletion at tumor initiation precludes SCLC-Aα development.

Differential transcriptomic analysis from the tumors in RPN and RP mice also revealed significantly downregulated genes in RPN tumors enriched for nervous system development (Figure 7E), suggesting that SCLC-Aσ is hierarchically closer to the SCLC-N and SCLC-Y/P subtypes, also supported by the data that variant morphology is more frequent in RPN. To address whether the absence of Nkx2-1 resulted in distinct tumor evolution in RPN, we profiled copy numbers of those mSCLCs and found that *Nfib* and *Mycl* loci are frequently amplified in RP model (6/10) consistent with previous reports; however, these amplifications were absent in all tumors in RPN model, suggesting the lack of Nkx2-1 predisposed those RPN tumors to undergo a distinct path leading to a different lineage class of tumors (Figure 7F and Figure E8C).

## DISCUSSION

In the current classification, SCLC-A constitutes 70% of SCLCs, while emerging evidence suggests its heterogeneity. Our study revealed distinctive epigenetic landscapes within SCLC-A that we designated SCLC-Aα and SCLC-Aσ, characterized by the presence of *NKX2-1* and *TCF4* SEs. NKX2-1 is required for the lung development and later specification of the peripheral lung, but also of bronchial NE cells (40). NKX2-1 is a well-known marker to diagnose LADCs (41), but it is also frequently expressed in SCLC. As NKX2-1 plays key roles in brain development (42–45), it has been speculated that NKX2-1 is involved in NE pathways, based on its expression enriched in SCLC-A (46) and its physical interaction with ASCL1 (29). In our study, while most SCLC-A expresses NKX2-1 at variable levels, only 48% had *NKX2-1* SE. Importantly, *NKX2-1* deletion in SCLC-Aσ and SCLC-N cells, where NKX2-1 protein is still expressed, had little cell growth effects, suggesting its relevancy only in SCLC-Aα.

Further, our data suggest this function is mediated through interaction with SOX1. Combinations of master TFs that assemble at tissue-specific cis-regulatory sites control distinct cellular functions essential for maintaining lineage states in many contexts. Therefore, while the role of SOX1 had never been described in the lung, it is plausible that their interaction is critical to retain unique NE state and it further suggests biological plasticity between glandular cell lineage and pulmonary NE lineage.

The roles of NKX2-1 in carcinogenesis are complex and remain poorly understood. Nkx2-1 deletion in neoplastic lungs causes loss of pulmonary identity and reversion to a foregut lineage. Our data suggest that Nkx2-1 contributes to maintaining a specific neural lineage state that have classical SCLC morphology. The observation that *Nkx2-1* deletion is tolerated at tumor initiation in the GEMM, but not in established human SCLC-Aα cell lines, provokes a few potential interpretations. First, the timing of the deletion might be critical in determining the outcome, as evidenced in other studies on lineage factors (47, 48). Second, as CMV-Cre targets multiple cell types, *Nkx2-1* deletion may favor origination from certain cell types, thus influencing tumor evolution, as reflected by the lack of amplification at *Nfib* and *Mycl.* Third, early/in situ RPN lesions that would have become SCLC-Aα might simply evolve to an alternative identity due to lack of *Nkx2-1.*

Together, we subdivided the major SCLC-A subtype into SCLC-Aα and SCLC-Aσ by revealing distinct epigenetic landscape in SCLC cell lines, human SCLC tumors, and GEMM. Collaborative NKX2-1/SOX1 regulation defines the unique lineage state for SCLC-Aα. This study will provide foundation for future identification of vulnerabilities and enable us to develop higher level of personalized therapeutic approaches to benefit SCLC patients.

## Acknowledgements

We thank Hong Cai, Yi Ge, Ying Liu, Peipei Guo, Yifei Sun, Xiaokun Liu, Korey Kam, Mingzhe Li, Megan Vetter, Tyler Jacks and Karen Yee for helpful discussions and technical assistance, Aleksandra Wroblewska and Brian D. Brown for providing a lentiviral vector for cloning sgRNAs; NextSeq Sequencing Facility of the Department of Oncological Sciences at Icahn School of Medicine at Mount Sinai (ISMMS), Saboor Hekmaty, Gayatri Panda and Ravi Sachidanandam for sequencing assistance. The authors also thank Taplin Mass Spectrometry Facility at Harvard Medical School, the Biorepository and Pathology Core Facility at ISMMS and Center for Comparative Medicine and Surgery at the Icahn School of Medicine at Mount Sinai. Graphical Abstract was created with BioRender.com.

