## Supplementary material for "Transcriptional circuitry of NKX2-1 and SOX1 defines an unrecognized lineage subtype of small cell lung cancer": Online Data Supplement

**Corresponding author:** Dr. Hideo Watanabe, Division of Pulmonary, Critical Care and Sleep Medicine, Department of Medicine, Icahn School of Medicine at Mount Sinai, New York, NY 10029, USA. Phone: 212-241-9242; Fax: 212-876-5519;.

#### SUPPLEMENTARY METHODS

##### *Cell Lines*

SCLC cell lines COR-L88, DMS 79, NCI-H187, NCI-H209, NCI-H69, NCI-H2171, SHP-77, HCC33, NCI-H82, NCI-H1963, NCI-H196, NCI-H841, NCI-H2081, NCI-H524 and LADC cell lines NCI-H3122, NCI-H2087 were cultured in RPMI 1640 growth medium supplemented with 10% Fetal Bovine Serum (FBS) and 1% penicillin/streptomycin (P/S). NCI-H345 was maintained in HITES medium (serum free) with 1% P/S. NCI-H1436 was maintained in HITES medium with 5% FBS and 1% P/S. HEK293T cells were maintained in DMEM with 10% FBS and 1% P/S (Gibco). Cells were authenticated by copy number variation analyses (Figure S1C) and regularly tested for mycoplasma using the mycoAlert Detection Kit (Lonza).

##### *Chromatin Immunoprecipitation followed by sequencing (ChIP-seq)*

Approximately 20 $\times 10^6$ cells were crosslinked with 1% formaldehyde in PBS for 12 min at room temperature, quenched with 5 mg/ml bovine serum albumin (BSA) in Phosphate-buffered saline (PBS) and then washed twice with cold PBS. Cell pellets were resuspended in 1ml lysis buffer (50 mM Tris-HCl, pH 8.1, 10 mM EDTA, 1% SDS, 1× protease inhibitor cocktail (Thermo Fisher Scientific)) and sonicated with Diagenode Bioruptor sonicator for 13 cycles to obtain chromatin fragment lengths of 100 to 1,000 bp evaluated by Bioanalyzer DNA High sensitivity kit (Agilent). Fragmented chromatin was diluted in IP buffer (20 mM Tris-HCl pH 8.1, 150 mM NaCl, 2 mM EDTA, 1% Triton X-100) and incubated overnight at 4 °C with Protein G magnetic beads (Dynabeads: Life Technologies) that had been pre-incubated with anti-H3K27ac (Abcam, ab4729) or anti-NKX2-1 (Bethyl Laboratories, A300-BL4000) or anti-SOX1 (R&D, AF3369) antibodies. Immunoprecipitants were washed six times with wash buffer (50 mM HEPES pH 7.6, 0.5 M LiCl,

1 mM EDTA, 0.7% Na deoxycholate, 1% NP-40) and twice with TE buffer. Immunoprecipitated and IP input was treated with RNase A and Proteinase K on the beads, recovered in 1% SDS and 0.1 M NaHCO3 over a period of 6 h at 65 °C, and purified with DNA clean and concentrator-25 (Zymo Research). Up to 10 ng of DNA was used for the library construction using NEBNext Ultra II DNA Library Prep Kit (NEB, E7645). Sequencing was performed on NextSeq 500 (Illumina) for 38 nucleotides from paired ends according to the manufacturer's instructions.

***ChIP-seq data analysis***

*ChIP*-seq analyses were performed as described previously (1). For NKX2-1 and SOX1 ChIP, peaks were identified by MACS (2) after aligning to hg19/GRCh37 with Bowtie2. ChIP signals on NKX2-1 and SOX1 peaks detected on each sample were visualized as a heatmap generated by plotHeatmap function of deepTools (3). Functional analysis of the differentially bound regions was performed using Genomic Regions Enrichment of Annotations Tool (GREAT) (4).

***RNA-seq***

Total RNA was extracted using the RNeasy kit (Qiagen) and poly adenylated RNAs were enriched using NEBNext PolyA mRNA Magnetic Isolation Module (NEB, E7490) from 1 ug of total RNA for each sample, fragmented by incubating at 94°C for 15 min. First and second strand cDNAs were synthesized using SuperScript III reverse transcriptase (Thermo Fisher Scientific) and NEBNext® Ultra™ II Directional RNA Second Strand Synthesis Module (NEB). Sequencing library was generated using up to 10 ng of cDNA with NEBNext Ultra DNA Library Prep Kit (NEB, E7645). Paired-end sequencing was performed on NextSeq500 (Illumina) for 38 nucleotides from each end according to the manufacturer's instructions. The RNA-seq of RP and RPN mice tumors were performed differently as described previously (5).

***RNA-seq analysis***

RNA-seq analyses were performed as described previously (1). Briefly, DEseq2 (6) was used to identify the differentially expressed genes among control NCI-H187 cells (NCI-H187 parental cells and ones transduced with non-targeting sgRNAs) and NCI-H187 cells transduced with sgRNA targeting *NKX2-1* (sg1, sg2, sg3) and NCI-H187 sgRNA (sg1, sg2, sg3) targeting *SOX1*. Significantly differentially expressed genes were determined based on cutoffs of fold change > 1.4 and adjusted $P < 0.05$ for sg*NKX2-1* and fold change > 2 and adjusted $P < 0.05$ for sg*SOX1*. To identify potentially enriched functions of selected gene sets of interest, we compared these gene sets with the genes annotated by the Gene Ontology (GO) terms curated in the Molecular Signature Database (MSigDB) (7). Each of 7,481 GO terms included in "C5, GO biological process" collection was compared with query gene sets using Fisher's exact test. The significance of the overlap was determined based on p-value adjusted for multiple comparisons (FDR<0.05). Any GO terms consisting of more than 2,000 genes were considered non-specific and removed from the analysis.

***Lentiviral transduction of genes***

*FLAG-NKX2-1* or *FLAG-GFP* open reading frame (ORF) was cloned into pLEX_306 (a gift from David Root, Addgene plasmid #41391) using the Gateway cloning methods according to manufacturer's recommendations. HEK293T cells were seeded in a 10cm tissue culture dish and incubated at 37°C and 5% $CO_2$. At 80% confluency the cells were co-transfected with 10 μg of plasmid constructs, 7.5 μg of psPAX2 (a gift from Didier Trono, Addgene #12260) and 2.5 μg of pMD2.G (a gift from Didier Trono, Addgene #12259) vectors using TransIT-Lenti (Mirus) following manufacturer's recommendations. At 48h post-transfection, virus-containing supernatants were harvested, filtered (0.45 μm) and stored at −80°C. Cells were infected with lentiviral supplemented with polybrene at a final concentration of 8 μg/mL or lentiBlast (OZ BioSciences) at a ratio of 1:1000. Cells were selected with puromycin (1-2 μg/mL for 6 days).

***CRISPR-Cas9 genome editing***

Cells stably expressing Cas9 were generated by infection with the lentiCas9-Blast plasmid (Addgene # 52962, a gift from Feng Zhang). sgRNAs targeting *NKX2-1* or *SOX1* were selected from Brunello library (8). Non-target sgRNAs from the Gecko library v2 (9) which were used as scramble sgRNAs are listed in Table S1. sgRNAs were cloned using BbsI site downstream of the human U6 promoter in a lentiviral vector containing eGFP downstream of the human PGK promoter (a kind gift from Brown laboratory, ISMMS). Lentivirus was produced as above. Cells were first infected with the lentiCas9-Blast, and then selected with blasticidin (5μg/mL for 14 days), then infected with respective pLenti-GFP-sgRNA.

**Im*munohistochemistry (IHC)***

Immunohistochemical analyses were performed on xenograft tumor specimens, human primary SCLC tumor specimens and genetic engineered mouse model SCLC tumors. Tissues were embedded in paraffin following standard processing with ethanol dehydration. Tissue sections (5 μm thick) were deparaffinized and hydrated by standard methods, then heated at 95°C in 10 mM citrate buffer (pH 6.0) for antigen retrieval for 30 minutes. Endogenous peroxidase was quenched with 0.3% $H_2O_2$ in Tris-buffered saline (TBS) for 15 minutes. Blocking was performed with 10% normal horse serum in TBS at room temperature for 30 minutes. The tissues were incubated with the anti-NKX2-1 (1:100; Santa Cruz, sc-53136), anti-SOX1 (R&D, AF3369) or anti-ASCL1 (1:100, BD, 556604) antibody at 4°C overnight, and then washed with TBS-T (5 minutes, twice), followed by incubation with biotinconjugated horse anti-mouse secondary antibody (Vector Laboratories) at room temperature for 1 hour. Antibody binding was detected with the Vectastain ABC kit and diaminobenzidine (DAB) as substrate. All slides were counterstained with hematoxylin before being mounting.

***Western Blot Analyses***

To determine protein expression, cell lysates were prepared by incubating cells in lysis buffer (150 mM NaCl, 50 mM Tris-HCl at pH 8.0, 1% NP-40, 0.5% Na deoxycholate, 0.1% SDS, protease & phosphatase inhibitors) for 30 min at 4°C. After centrifugation to remove insoluble debris, protein concentrations were quantified using Pierce BCA (Thermo Fisher Scientific). Proteins were separated by electrophoresis on a SDS-PAGE gel (BioRad), transferred to a PVDF membrane (Thermo Fisher Scientific) and blocked with 5% milk in TBS-T. The membranes were immunoblotted with anti-NKX2-1 antibody (Bethyl Laboratories, A300-BL4000), anti-SOX1 (R&D, AF3369), anti-ASCL1 (BD, 556604) or anti-vinculin (Sigma).

***Co-immunoprecipitation***

Stable NCI-H187, NCI-H209, or NCI-H1436 cells expressing pLEX_306_FLAG-NKX2-1 or pLEX_306_FLAG-GFP were established by infecting with lentivirus using standard viral transfer protocol followed by puromycin selection. Chromatin fraction was prepared from these stables cells and subsequently co-immunoprecipitated with the methods described previously (10) with a minor modifications. Briefly, the nuclear pellets were lysed in 5 volumes of FLAG-IP buffer (150 mM NaCl, 50 mM Tris pH7.5, 1 mM EDTA, 0.5% NP40, 10% glycerol, protease inhibitors) for 30 min. The insoluble pellet was washed in MNase buffer (50 mM Tris-HCl (pH7.9), 5 mM $CaCl_2$). The pellet then was thoroughly resuspended in five pellet volume of MNase buffer containing 2.5ul of Micrococcal nuclease (NEB, M0247S), incubate for 20 min at 37°C with frequent mixing.

Stop the digest by adding 0.5 M EDTA, then diluted with an equal volume of 2x IP high salt buffer (600 mM NaCl, 50 mM Tris (pH 7.5), 2 mM EDTA, 1% NP40, 20% Glycerol, Protease inhibitors). The supernatant containing the solubilized chromatin extract was then incubated with Protein G magnetic beads (Dynabeads: Life Technologies) that had been pre-incubated with anti-FLAG M2 (Sigma, F1804) antibody overnight, followed by elution with SDS-PAGE sample buffer. The final elution was further processed for IP-WB or LC/MS/MS sequencing and data analysis.

***Protein Sequence Analysis by LC-MS/MS***

Excised gel bands were cut into approximately 1 $mm^3$ pieces. Gel pieces were then subjected to a modified in-gel trypsin digestion procedure. Gel pieces were washed and dehydrated with acetonitrile for 10 min. followed by removal of acetonitrile. Pieces were then completely dried in a speed-vac. Rehydration of the gel pieces was with 50 mM ammonium bicarbonate solution containing 12.5 ng/µl modified sequencing-grade trypsin (Promega, Madison, WI) at 4ºC. After 45 min., the excess trypsin solution was removed and replaced with 50 mM ammonium bicarbonate solution to just cover the gel pieces. Samples were then placed in a 37ºC room overnight. Peptides were later extracted by removing the ammonium bicarbonate solution, followed by one wash with a solution containing 50% acetonitrile and 1% formic acid. The extracts were then dried in a speed-vac (~1 hr). The samples were then stored at 4ºC until analysis. On the day of analysis, the samples were reconstituted in 5 - 10 µl of HPLC solvent A (2.5% acetonitrile, 0.1% formic acid). A nano-scale reverse-phase HPLC capillary column was created by packing 2.6 µm C18 spherical silica beads into a fused silica capillary (100 µm inner diameter x ~30 cm length) with a flame-drawn tip. After equilibrating the column each sample was loaded via a Famos auto sampler (LC Packings, San Francisco CA) onto the column. A gradient was formed, and peptides were eluted with increasing concentrations of solvent B (97.5% acetonitrile,

0.1% formic acid). As peptides eluted, they were subjected to electrospray ionization and then entered into an LTQ Orbitrap Velos Pro ion-trap mass spectrometer (Thermo Fisher Scientific, Waltham, MA). Peptides were detected, isolated, and fragmented to produce a tandem mass spectrum of specific fragment ions for each peptide. Peptide sequences (and hence protein identity) were determined by matching protein databases with the acquired fragmentation pattern by the software program, Sequest (Thermo Fisher Scientific, Waltham, MA). All databases include a reversed version of all the sequences and the data was filtered to between a one and two percent peptide false discovery rate.

***Cell proliferation assay***

Cells were plated at a density of 3,000 cells/well with five replicates in a 96-well plate; five identical plates were prepared. Cell viability was assayed at 1, 3, 5, 7 and 9 days after plating with alamarBlue Cell Viability Reagent (Thermo Fisher) and fluorescence at 585 nm was measured on a Spectra Max3 plate reader (Molecular Device) according to the manufacturer's protocol at excitation of 555 nm. Cell viability at 3, 5, 7 and 9 days were corrected for the ratio to control cells from the day 1 reading to account for plating unevenness.

***Xenograft model***

NCI-H187 and NCI-H209 cell lines expressing non-target sgRNA or NKX2-1 sgRNA were cultured with RPMI 1640 medium. Cells ($5 \times 10^6$) were injected with a 1:1 mixture of 50 µl cell suspension and 50 µl Matrigel (Corning) subcutaneously into both flank regions of 4- to 6-week old male NOD-scid gamma mice (Jackson Laboratory). Tumor volume (length x width$^2$ /2) was measured twice a week. When the largest tumor size reached at 1000 mm$^3$, mice were sacrificed, and tumors were immersed in formalin for immunohistological analysis. All xenograft studies were conducted following approval from the institutional Animal Care and Use Committee at Icahn School of Medicine at Mount Sinai.

***Annexin-V staining***

Cells were harvested at four days after transduction and washed with PBS. The cells were stained with Annexin-V and propidium iodide using Alexa Fluor® 488 annexin V/Dead Cell Apoptosis Kit (Thermo Fisher Scientific) according to manufacturers' recommendations. Fluorescence was measured by FACS Canto II (BD Bioscience). Quantitative analyses for the cell viability proportions were performed using FlowJo software.

***In situ proximity ligation assay (in situ PLA)***

DMS 79, NCI-H187, NCI-H209, NCI-H345, or NCI-H841 cells were seeded on poly-L-Lysine coated coverslips. After 24 h, cells were fixed in 4% paraformaldehyde (PFA) for 10 min at 4°C, then permeabilized using 0.1% Triton X-100 for 10 min at room temperature. *In situ* PLA was performed with DuoLink In Situ Reagents (Sigma) according to the manufacturer's instructions. Primary antibodies used were: rabbit anti-NKX2-1 (Bethyl Laboratories, A300-BL4000), goat anti-SOX1 (R&D, AF3369), rabbit isotype control (Cell signaling, 3900s), and normal goat IgG control (R&D, AB-108-C). Images were acquired with a ZEISS Axiocam 503 mono confocal microscope.

***Super-Enhancer analysis***

Super-enhancers were defined using the Rank Ordering of Super-Enhancers (ROSE2) algorithm (11). H3K27ac-enriched peaks were stitched together, using an optimal distance determined by the algorithm per sample up to 12.5 kb. The algorithm generates plots for all stitched enhancers per each sample and defines super-enhancers on the basis of top-ranked enhancers with highest read counts at the tangential cutoff. Unsupervised hierarchical clustering was performed using signals (read counts on defined super-enhancer regions) near transcriptional regulators defined by the term GO0003677 (DNA-binding).

***Transcriptional Regulatory Networks***

Core transcriptional regulatory circuitries (CRCs) were determined by CRCmapper (12) from SE determined as above at the default parameters CRCs in all 25 SCLC cell lines. Subtype specific common CRCs were determined by the frequency of the factor in the CRCs (>50%) in the subtype, and the proportion of samples with the factor in CRCs that belong to the subtype (>50%).

***Fixed-tissue ChIP-seq for H3K27 acetylation (H3K27ac) (FiTAc-seq) for human and mice tumors***

FiTAc-seq has been previously described (13), briefly, ten 1mm mice tumor tissue cores or two sections of 50um thickness for full sections of human FFPE tumors were used for ChIP profiling. Samples were treated with xylene to remove paraffin and then rehydrated with 100% ethanol for 10 min, 70% ethanol for 10 min, and water for another 10 min. Hydrated tissues were resuspended in 200ul of 1% SDS lysis buffer with 4ul of 50× EDTA-free protease inhibitor cocktail (PIC) and 1ul of 1 M sodium butyrate, then incubated overnight at 65°C with constant agitation at 1,000 r.p.m. The homogenized tissues were sonicated with Bioruptor for 12 cycles of 30 second on and 30 second off at 4°C. The soluble chromatins were subject to ChIP assay as described above. Library preparation and sequencing were done as described above.

***Copy number analysis***

Mouse tumors copy number analysis of somatic alterations was performed using ChIP-seq input data (effectively low-pass whole genome sequencing data) on tumors from RP or RPN. Copy numbers were inferred and visualized using CNVkit (14).

***Mice and tumor initiation***

Mice harboring $Rb1^{flox}$, $Trp53^{flox}$ and $Nkx2\text{-}1^{flox}$ alleles have been previously described (15, 16). Tumors were generated by intratracheal delivery of Ad5CMV-Cre adenovirus (University of Iowa, Gene Transfer Vector Core) as described (17). Viral dose was 5x107 plaque forming units (pfu)/mouse unless otherwise specified. Animal studies were approved by the Committee for Animal Care at MIT, and conducted in compliance with the Animal Welfare Act Regulations and other federal statutes relating to animals and experiments involving animals and adheres to the principles set forth in the Guide for the Care and Use of Laboratory Animals, National Research Council, 1996 (institutional animal welfare assurance no. A-3125–01).

#### SUPPLEMENTARY TABLE

**Table E1:** Cell lines used in this study

| Cell lines | Source | Identifier |
|---|---|---|
| Human: HEK-293T | ATCC | ATCC Cat# CRL-3216, RRID:CVCL_0063 |
| Human: DMS 79 | ATCC | ATCC Cat# CRL-2049, RRID:CVCL_1178 |
| Human: NCI-H187 | Gift of Israel Cañadas, Dana-Farber Cancer Institute | ATCC Cat# CRL-5804, RRID:CVCL_1501 |
| Human: NCI-H209 | ATCC | ATCC Cat# HTB-172, RRID:CVCL_1525 |
| Human: NCI-H345 | Gift of Israel Cañadas, Dana-Farber Cancer Institute | ATCC Cat# HTB-180, RRID:CVCL_1558 |
| Human: NCI-H1436 | Gift of Israel Cañadas, Dana-Farber Cancer Institute | ATCC Cat# CRL-5871, RRID:CVCL_1471 |
| Human: NCI-H196 | Gift of Israel Cañadas, Dana-Farber Cancer Institute | ATCC Cat# CRL-5823, RRID:CVCL_1509 |
| Human: NCI-H841 | Gift of Israel Cañadas, Dana-Farber Cancer Institute | ATCC Cat# CRL-5845, RRID:CVCL_1595 |
| Human: NCI-H2081 | Gift of Israel Cañadas, Dana-Farber Cancer Institute | ATCC Cat# CRL-5920, RRID:CVCL_1522 |
| Human: NCI-H524 | Gift of Israel Cañadas, Dana-Farber Cancer Institute | ATCC Cat# CRL-5831, RRID:CVCL_1568 |
| Human: NCI-H1963 | ATCC | ATCC Cat# CRL-5982, RRID:CVCL_1510 |
| Human: NCI-H82 | Gift of Israel Cañadas, Dana-Farber Cancer Institute | ATCC Cat# HTB-175, RRID:CVCL_159 |
| Human: HCC33 | ATCC | DSMZ Cat# ACC-487, RRID:CVCL_2058 |
| Human: SHP-77 | Gift of Israel Cañadas, Dana-Farber Cancer Institute | ATCC Cat# CRL-2195, RRID:CVCL_1693 |
| Human: NCI-H2171 | ATCC | ATCC Cat# CRL-5929, RRID:CVCL_1536 |
| Human: NCI-H69 | ATCC | ATCC Cat# HTB-119, RRID:CVCL_1579 |
| Human: COR-L88 | Gift of Israel Cañadas, Dana-Farber Cancer Institute | ECACC Cat# 92031917, RRID:CVCL_1141 |

**Table E2: sgRNA sequences for the CRISPR/Cas9 system**

| | sgRNA sequence (5’-3’) | PAM sequence |
|---|---|---|
| NKX2-1_sg1 | GCGAGCGGCATGAACATGAG | CGG |
| NKX2-1_sg2 | CAAGCAACAGAAGTACCTGT | CGG |
| NKX2-1_sg3 | CGCCGCCTACCACATGACGG | CGG |
| SOX1_sg1 | CAAGAAGGACAAGTACTCGC | TGG |
| SOX1_sg2 | CCCATGCACCGCTACGACAT | GGG |
| SOX1_sg3 | CGAGTGGAAGGTCATGTCCG | AGG |
| non-target sg1 | ACGGAGGCTAAGCGTCGCAA | N/A |
| non-target sg2 | CGCTTCCGCGGCCCGTTCAA | N/A |

#### SUPPLEMENTARY FIGURES

**Figure E1 related to Figure 1: Epigenomic profiling subdivided SCLC-A by differential super-enhancers**

**A.** PCA showing the distinction of SE signals near transcriptional regulator genes in each cell line among four SCLC Clusters. **B.** Genome view tracks of H3K27ac signal at the *ASCL1 and TCF4* loci in four clusters of SCLC cell lines. **C.** Copy number profiles showing genes amplification in chromosome 14 in 4 SCLC-Aα (COR-L88, DMS 79, NCI-H187 and NCI-H209) and 3 LADC (NCI-H2087, NCI-H3122 and NCI-H441). *NKX2-1* locus is highlighted in grey box. **D.** SE plots with all enhancers rank-ordered by H3K27ac signals in COR-L88 (SCLC-Aα) and NCI-H146, SHP-77, NCI-H69 and NCI-H2081 (SCLC-Aσ) . Dotted lines depict tangential cutoffs to define SEs. **E.** H3K27ac ChIP-seq track of *NKX2-1* locus is shown in 4 SCLC-Aα (COR-L88, DMS 79, NCI-H187 and NCI-H209) and 2 LADC (NCI-H2087 and NCI-H3122). **F.** Genome view tracks of H3K27ac signal at the *ASCL1*, *NEUROD1*, *POU2F3*, *YAP1* and *NKX2-1* loci in 4 human SCLC specimens of 3 SCLC-N and 1 SCLC-P subtypes.

**Figure E2 related to Figure 1: Epigenomic profiling subdivided SCLC-A by differential super-enhancers**

**A.** Images from IHC analysis of consecutive sections on Tumor 1 for ASCL1, NEUROD1, NKX2-1 and H&E, Scale bars, 25μm. **B.** Unsupervised clustering of 135 primary tumors and 50 CCLE cell lines using the expression of *ASCL1*, *NEUROD1*, *POU2F3*, *YAP1* and *NKX2-1* distinguishing five subgroups. **C.** Box and jitter plots showing the distribution of *NKX2-1* mRNA expression in 135 primary SCLC tumors, along with that in LADC and LUSC tumors from TCGA. **D.** Ponceau S staining as a loading control for the Western blot panel in Figure 1F.

**Figure E3 related to Figure 2: *NKX2-1* survival dependency is unique to SCLC-Aα**

**A.** Fluorescence-activated cell sorting (FACS) analysis of annexin $V^+$ cells in NT g1 and NKX2-1 sg1 of NCI-H187 cells. **B.** Tumor growth curve for xenograft tumors of NCI-H187 cells infected with *NKX2-1* sg3 or NT sg2 *in vivo*. Mean ± SD of four biological replicates. *$P < 0.0001$, t test. Xenograft tumors were resected at 3 weeks after inoculation. **C.** H&E staining and IHC staining of NKX2-1 and GFP in NCI-H187 NT sg1 and NCI-H187 NKX2-1 sg1 xenograft tumors. Scale bars, 50μm. **D.** Immunoblots showing protein expression of NKX2-1, ASCL1 and vinculin as a loading control in SCLC-Aα cell lines NCI-H187 and NCI-H209 with three *NKX2-1* sgRNAs, or two non-target (NT) sgRNAs.

**Figure E4 related to Figure 3: Genomic occupancy of NKX2-1 in SCLC-Aα cells distinct from that in LADC cells**

**A.** PCA of NKX2-1 bound sites in 5 SCLC-Aα cell lines and 3 LUAC cell lines. Each dot represents a cell line. **B.** Volcano plot showing 2,671 differentially NKX2-1 bound loci (661 enriched in SCLC-Aα and 2,010 enriched in LADC) at thresholds of absolute fold change $\geq 2^5$ and FDR $\leq 1x10^{-5}$ in 5 SCLC-Aα cell lines and 3 LUAC cell lines. **C.** Enriched ontology by GREAT analyses for regions differentially bound in 3 LUAC cell lines. **D.** Motif-based analysis of sequences from the differential NKX2-1-binding in 3 LUAC cell lines and 5 SCLC-Aα cell lines.

**Figure E5 related to Figure 4: NKX2-1 interactome in chromatin of SCLC-Aα cells includes SOX1**

**A.** Protein expression of NKX2-1 and vinculin as a loading control in NCI-H187 and NCI-H1436 parental, FLAG-GFP and FLAG-NKX2-1 cells. **B.** CRCs commonly identified in 6 SCLC-Aσ cell lines, 5 SCLC-N cell lines, 6 SCLC-P/Y cell lines. **C.** Genome view tracks of H3K27ac signal at the *NFIB, PBX1* and *TCF12* loci in four clusters of SCLC cell lines. **D.** Box and jitter plots showing the distribution of *SOX1* mRNA expression in SCLC-A, SCLC-N, SCLC-P, SCLC-Y subtypes of 135 primary SCLC tumors. **E.** Correlation between *NKX2-1* and *SOX1* mRNA expression in 135 primary SCLC tumors and 50 CCLE cell lines. Each dot represents a tumor or a cell line. **F.** Protein expression of SOX1, NKX2-1, ASCL1 and vinculin as a loading control in SCLC cell lines from different clusters. *NKX2-1*-amplified adenocarcinoma cell lines NCI-H3122 and NCI-H2087 are shown as reference. **G.** In situ proximity ligation assay signals (red fluorescent dots) targeting endogenous SOX1 and NKX2-1 in NCI-H345 cells. Negative control NCI-H187 without NKX2-1/SOX1 primary antibodies is also shown. Nuclei are counterstained with DAPI (blue).

**Figure E6 related to Figure 5: NKX2-1 and SOX1 co-occupy the SCLC-Aα genome to collaborate in neuronal gene regulation**

**A.** Venn diagram showing NKX2-1 binding sites and SOX1 binding sites in NCI-H187, DMS 79, NCI-H209 and NCI-H345 cell lines. **B.** Enriched ontology by GREAT analyses for regions in 2,652 overlapping NKX2-1-SOX1 binding sites in NCI-H187 cell line.

**Figure E7 related to Figure 6: Collaboration of NKX2-1 and SOX1 is required to control the neuronal state in SCLC-Aα**

**A.** *(Top)* Immunoblots showing protein expression of SOX1, NKX2-1 and vinculin as a loading control in SCLC-Aα cell lines NCI-H187 and NCI-H209 with three *SOX1* sgRNAs, or two non-target (NT) sgRNAs. *(bottom)* Cell growth curve for SCLC-Aα cell lines infected with *SOX1* sgRNAs or NT sgRNAs. Data are shown as mean ± SD (n=3); *P < 0.0001 by two-sided two-way

ANOVA. **B.** Bar plots showing percentage of apoptotic NCI-H187 cells infected with NT sgRNA and *SOX1* sgRNA, measured by annexin V staining. Means ± SD of three biological replicates. *P < 0.0001, t test. Representative data for *NT* sg2 and *SOX1* sg2 are shown. **C.** PCA of NKX2-1 bound sites in NCI-H187 with three *SOX1* sgRNAs, two non-target (NT) sgRNAs and one parental cell. **D.** Volcano plot showing 2,067 differentially NKX2-1 bound loci signal reduced at thresholds of absolute fold change ≥ 2 and FDR ≤ 0.1 in 3 NCI-H187 *SOX1* sgRNAs cells compared to 3 control cells.

**Figure E8 related to Figure 7: *Nkx2-1* is essential for development of SCLC-Aα tumors in an autochthonous mouse model**

**A.** Genome view tracks at the *Neurod1*, *Yap1*, and *Pou2f3* loci for H3K27ac ChIP-seq signals in 10 SCLC tumors from RP mice and 10 SCLC tumors from RPN mice. **B.** Survival analysis of RP (n=14) and RPN (n=10) mice. **C.** Copy number profiles showing genes amplification of the *Nfib* and *Mycl* loci in chromosome 4 for 10 SCLC tumors each from RP and RPN mice.

Supplementary Figure 1

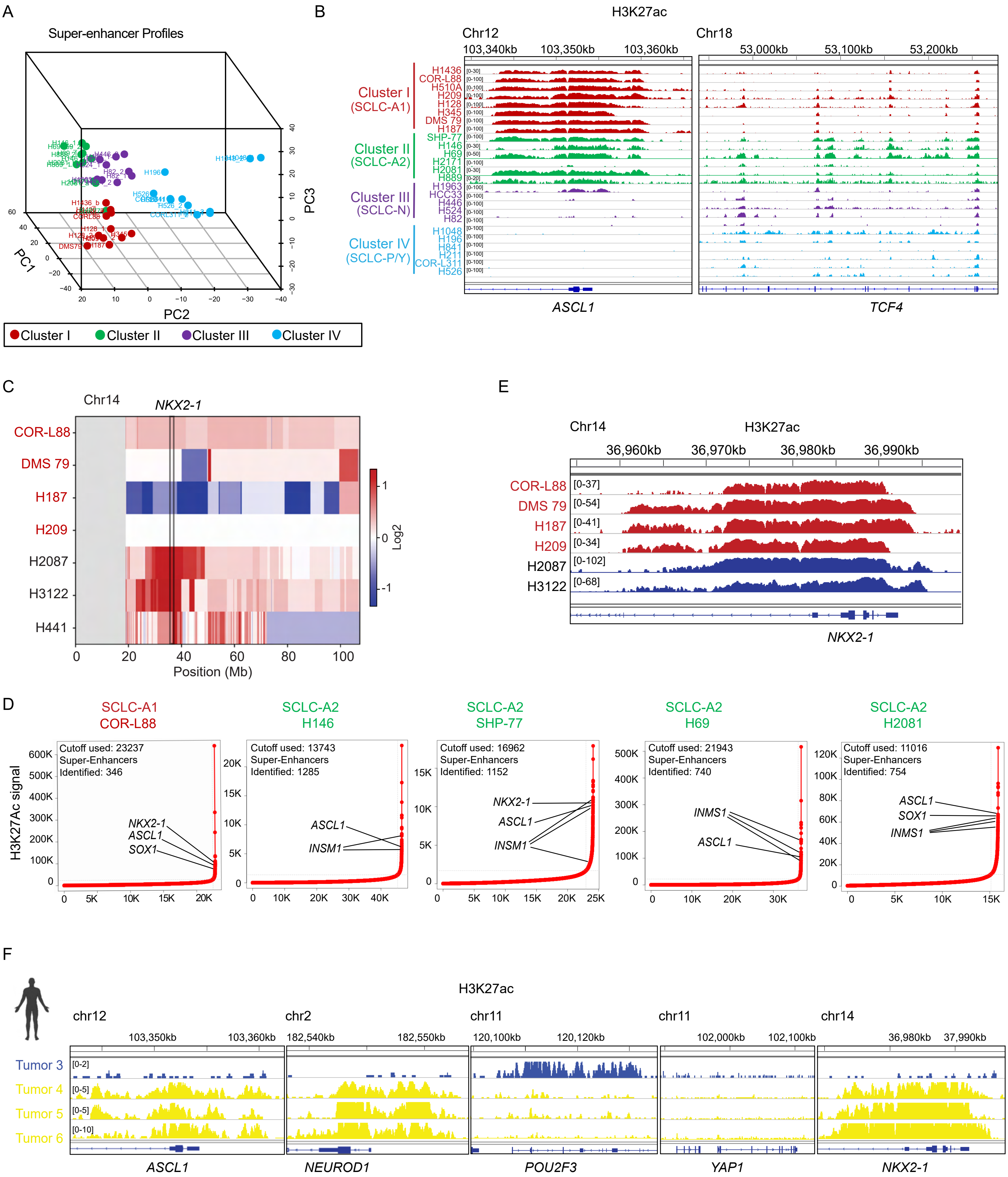

Supplementary Figure 2

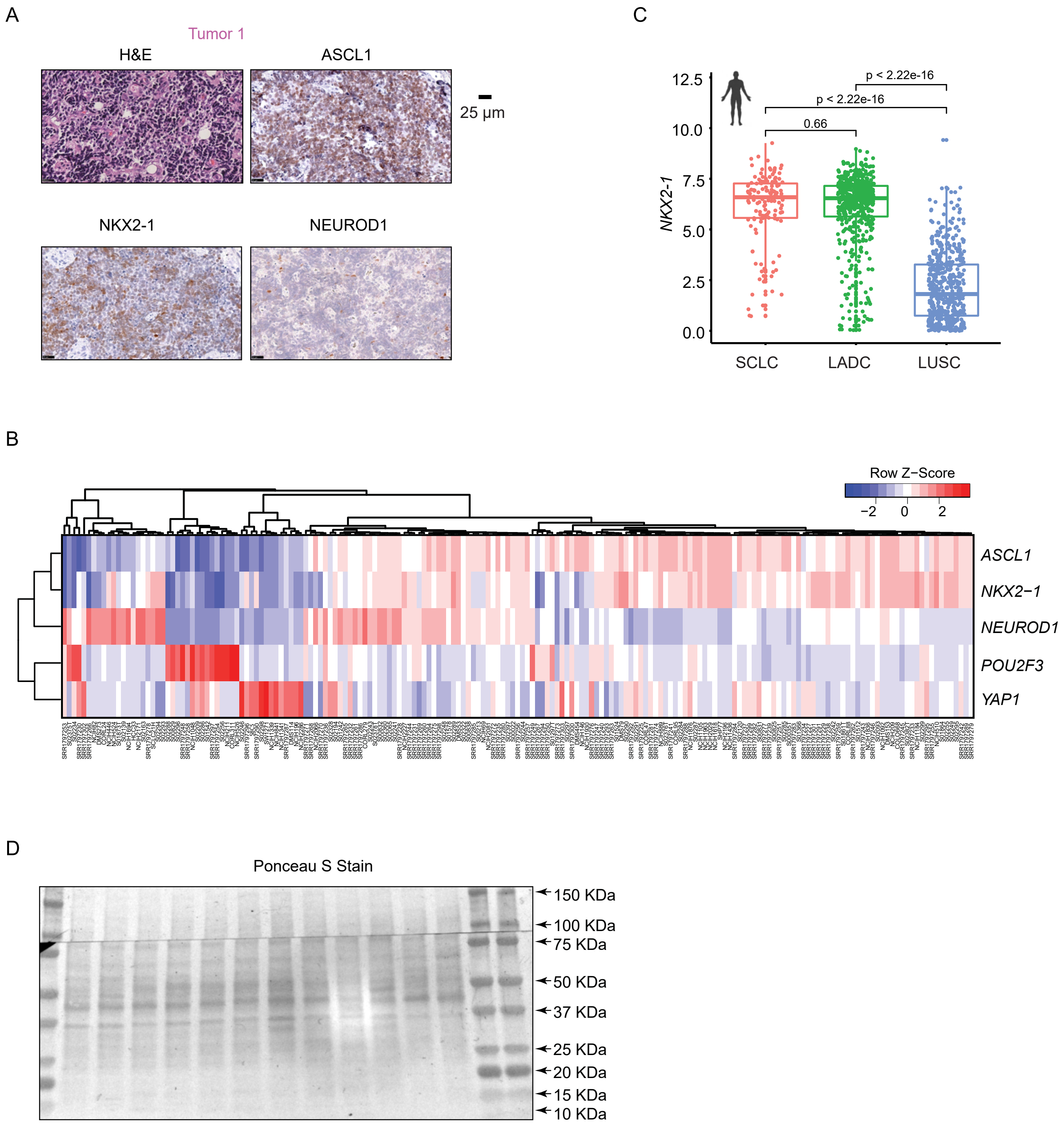

Supplementary Figure 3

A

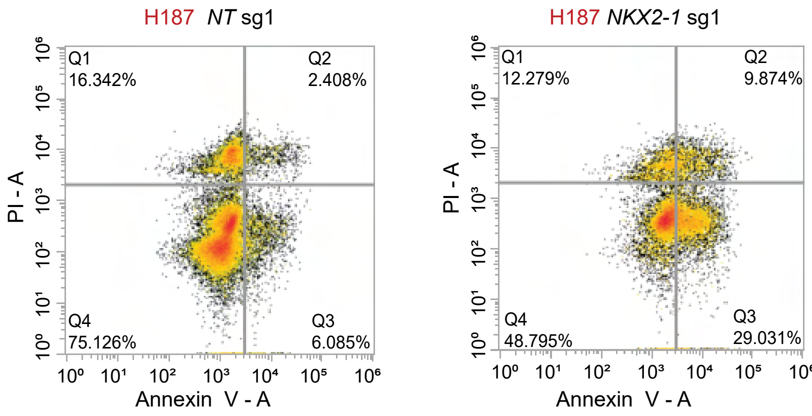


B

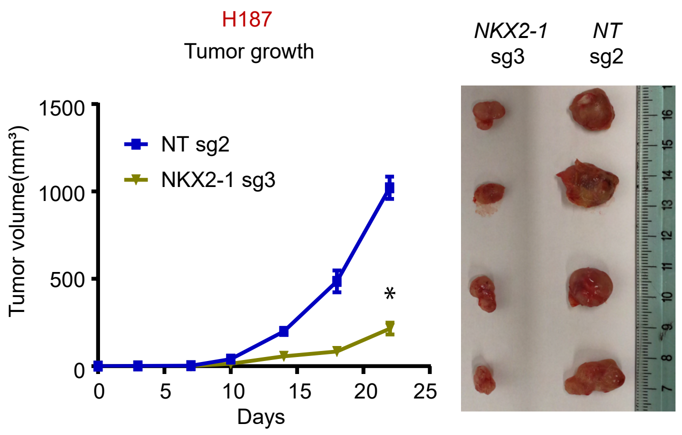


C

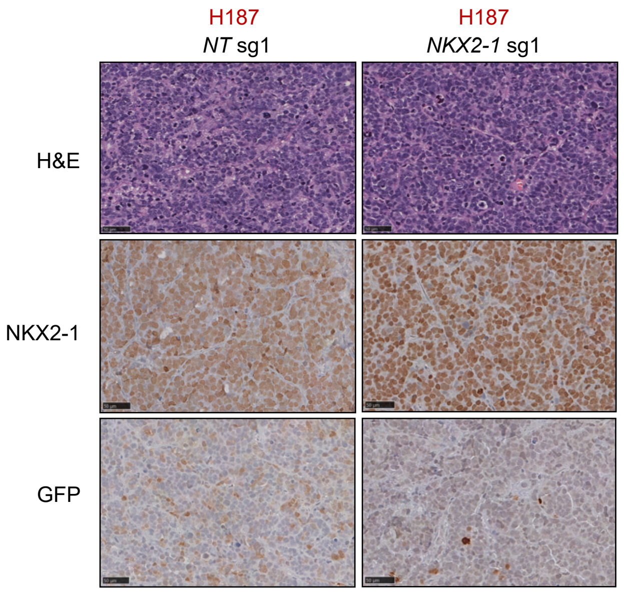


D

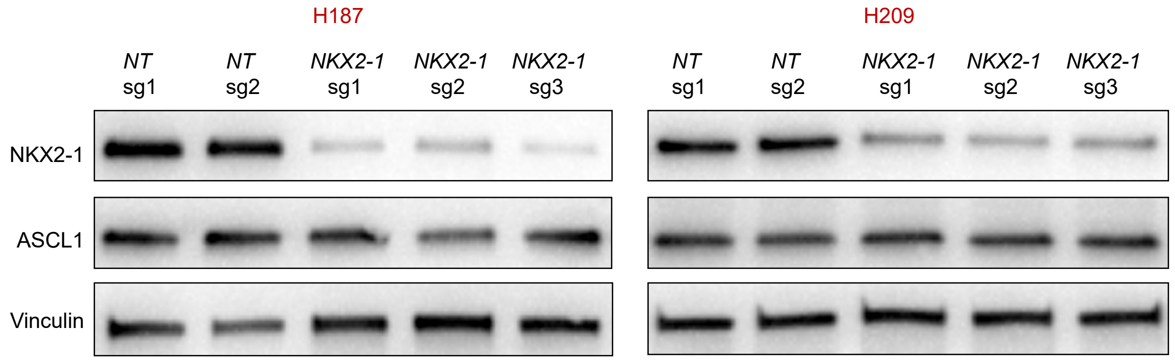

### Supplementary Figure 4

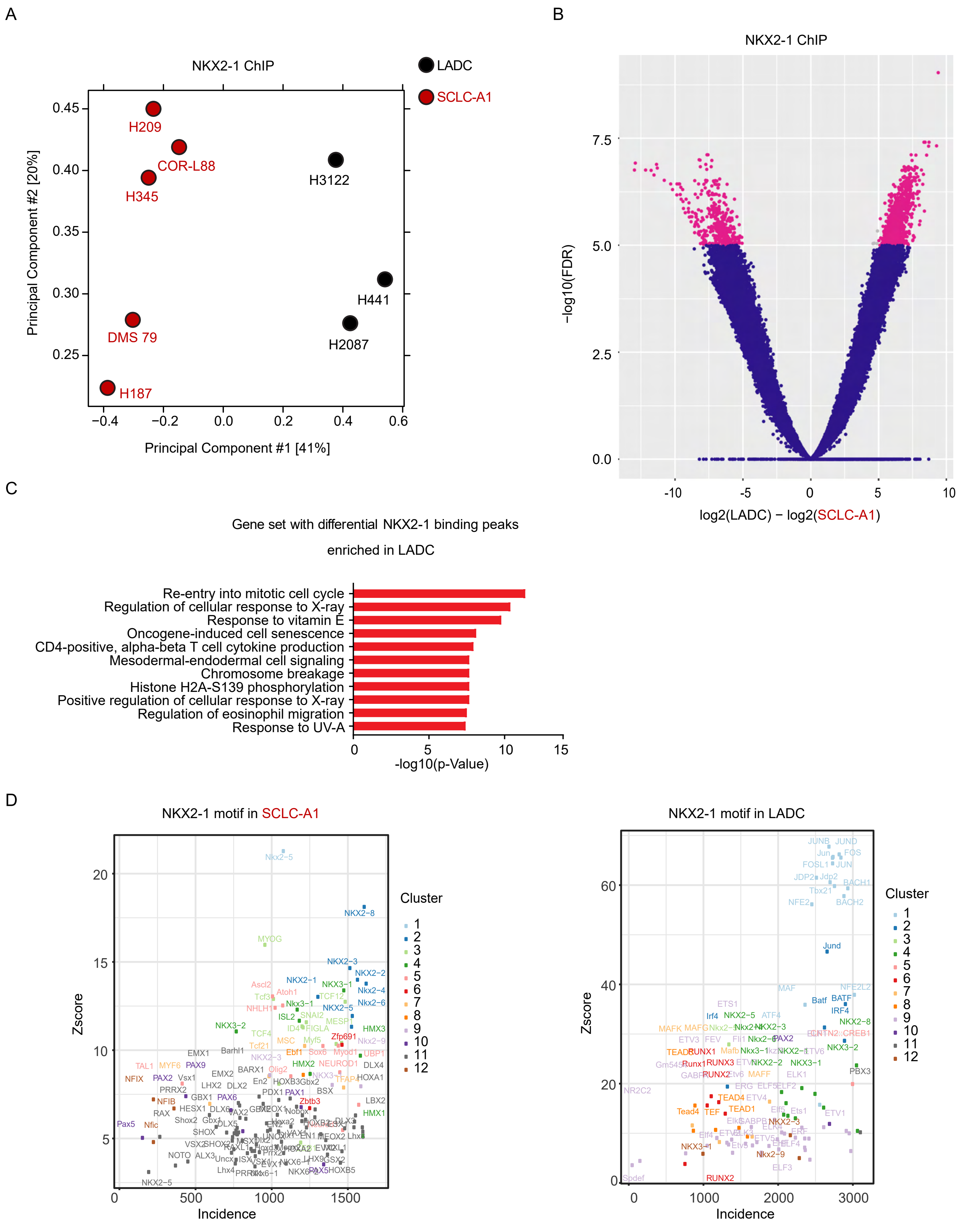

Supplementary Figure 5

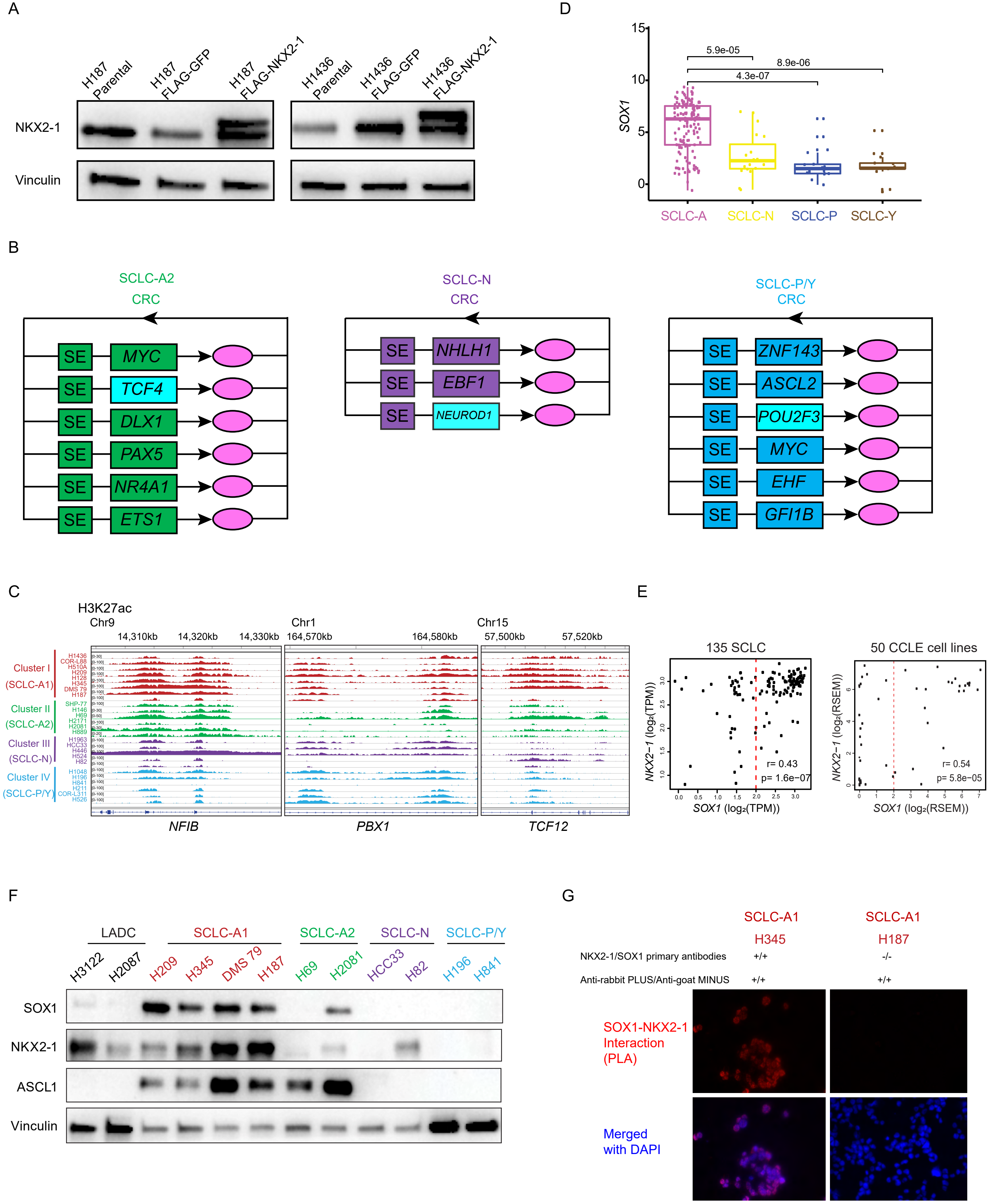

Supplementary Figure 6

A

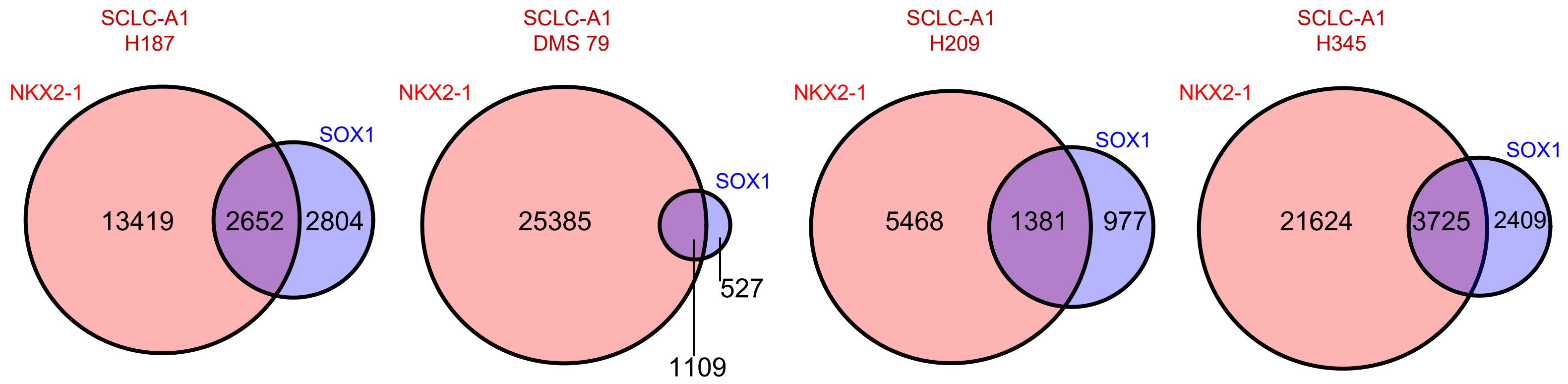


B

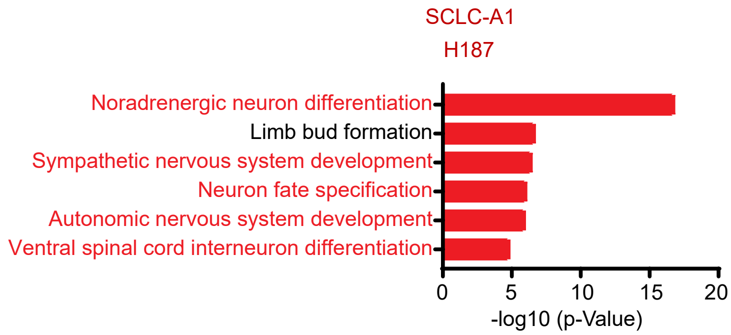

Supplementary Figure 7

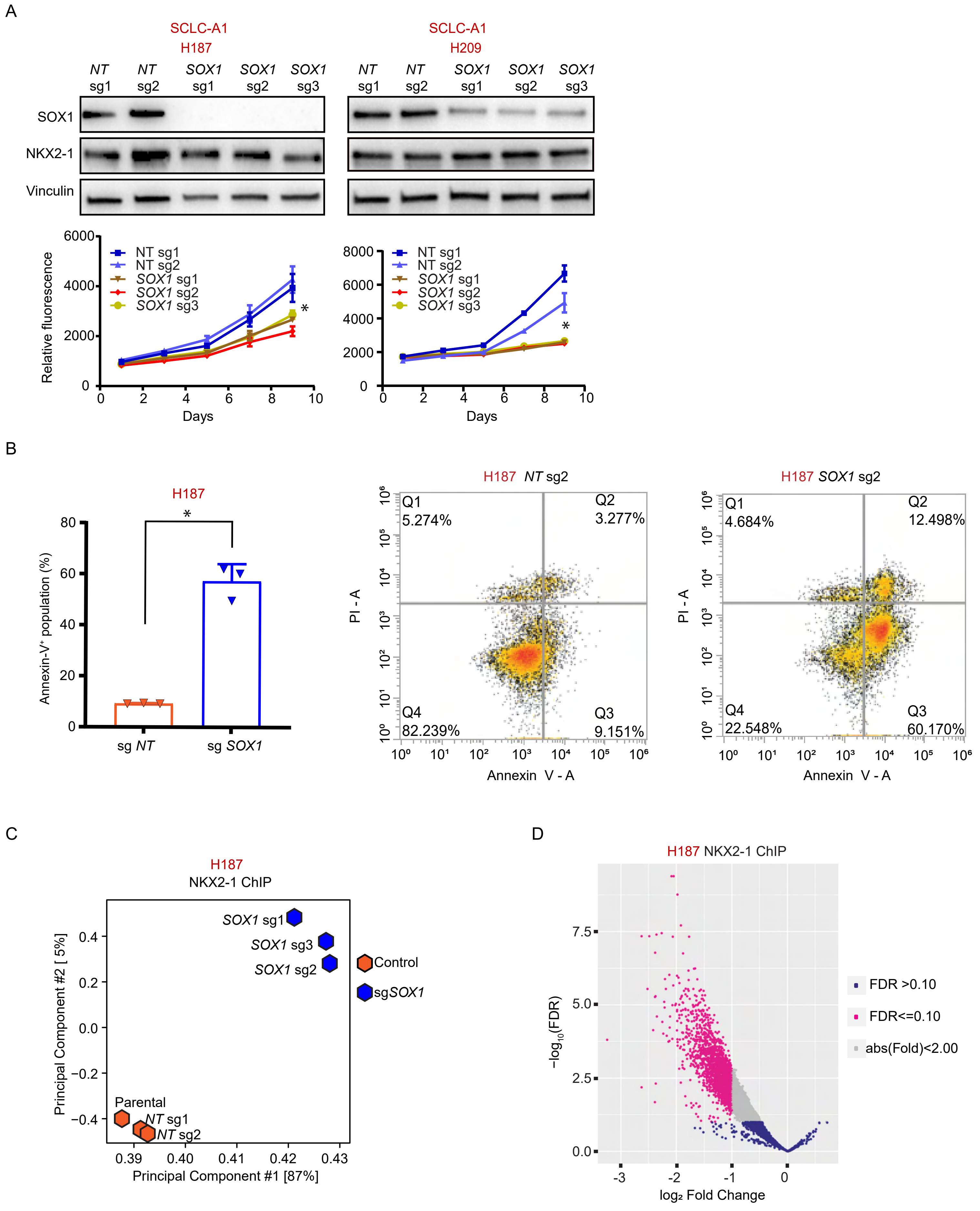

Supplementary Figure 8

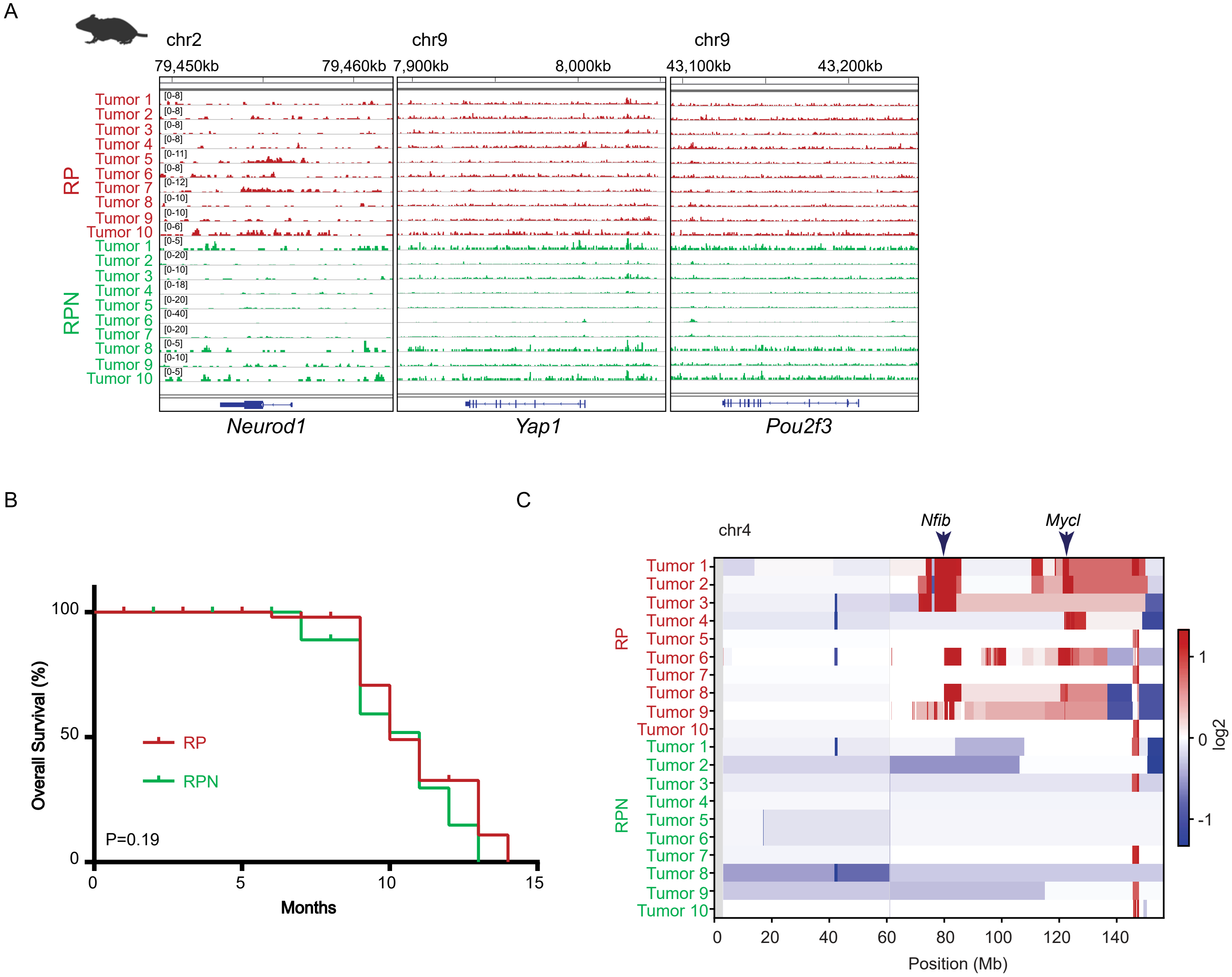
